# Microbial extracellular polysaccharide production and aggregate stability controlled by Switchgrass (*Panicum virgatum*) root biomass and soil water potential

**DOI:** 10.1101/724195

**Authors:** Yonatan Sher, Nameer R. Baker, Don Herman, Christina Fossum, Lauren Hale, Xing-Xu Zhang, Erin Nuccio, Malay Saha, Jizhong Zhou, Jennifer Pett-Ridge, Mary Firestone

**Affiliations:** University of California, Berkeley, CA; USDA-Agricultural Research Service, Parlier, CA; Lanzhou University, Lanzhou, China; Lawrence Livermore National Laboratory, Livermore, CA; Noble Research Institute, Ardmore, OK; University of Oklahoma, Norman, OK; Lawrence Berkeley National Laboratory, Berkeley, CA

**Author notes:** equal contribution (Yonatan Sher, Nameer R. Baker).

**Keywords:** EPS, marginal soil, Stable soil aggregate, switchgrass, ^13^C labeling

## Abstract

Deep-rooting perennial grasses are promising feedstock for biofuel production, especially in marginal soils lacking organic material, nutrients, and/or that experience significant water stress. Perennial grass roots can alter surrounding soil conditions and influence microbial activities, particularly the production of extracellular polymeric substances composed primarily of extracellular polysaccharides (EPS). These polymers can alleviate cellular moisture and nutrient stress, and enhance soil characteristics through improved water retention and aggregate stability, the latter of which may in turn enhance carbon persistence. In this study we used a ^13^CO_2_ tracer greenhouse experiment to examine the effect of switchgrass cultivation on the production and origin of EPS in a marginal soil with five fertilization/water treatments (control, +N, +NP, +P, low water). Soils with both added nitrogen and phosphorus had the highest root biomass, EPS and percentage of water-stable soil aggregates. Multiple linear regression analyses revealed root biomass was the most important determinant for soil EPS production, potentially by controlling carbon supply and diurnal changes in soil water potential. Path analysis highlighted the role of soil water potential were and EPS on with water-stable soil aggregates, indicating that EPS concentration and soil aggregation have similar drivers in this soil. High mannose content confirmed the microbial origin of EPS. ^13^CO_2_ labeling indicated that 0.18% of newly fixed plant carbon was incorporated into EPS. Analysis of field samples suggests that EPS is significantly enhanced under long-term switchgrass cultivation. Our results demonstrate that switchgrass cultivation can promote microbial production of EPS, providing a mechanism to enhance sustainability of marginal soils.

## 1. INTRODUCTION

In oligotrophic and variable moisture environments, such as those often found in marginal land soils, microorganisms can produce polymeric substances protecting them from external stresses (Cheshire, 1977; Nicolaus et al., 2010; Oades, 1984; Sandhya and Ali, 2015; Wolfaardt et al., 1999). These polymeric substances include a variety of biological polymers, such as DNA and proteins, but it has been shown that the principal components are polysaccharides (Cheshire, 1977; Hall-Stoodley et al., 2004; More et al., 2014; Oades, 1984); hence we will focus on extracellular polysaccharides (EPS). Physical and chemical characteristics of EPS can help microbial cells alleviate moisture and nutrient stress, and coincidently affect soil characteristics in a manner that may enhance the sustainability of marginal soils. EPS can mitigate the effect of decreasing water potential on microbial cells by increasing the water content they perceive (Adessi et al., 2018; Chenu, 1993; Sandhya and Ali, 2015). Enhanced soil water holding capacity can increase nutrient diffusion to and from microbial cells encased in EPS (Chenu and Roberson, 1996). The sticky, gelatinous properties of EPS also bind microbial cells to mineral surfaces in soil (Wolfaardt et al., 1999) and enhances soil-aggregation by binding soil particles together (Costa et al., 2018; Oades, 1984; Rogers and Burns, 1994) into micro-aggregates (Six et al., 2000; Six and Paustian, 2014). EPS thus enhances the formation of water-stable aggregates (Sandhya and Ali, 2015) and increases mean soil aggregate size (Amellal et al., 1999). EPS may also increase C persistence in soil, as it can be occluded in tertiary structures where it is less available for microbial consumption (Chenu and Plante, 2006; Liang et al., 2017).

Microbial EPS production tends to increase during dry periods, as microbes produce more EPS to enhance water retention of the surrounding soil matrix (Roberson and Firestone, 1992). As such, we would expect to see more EPS produced in drier or variable moisture soil conditions. EPS production is also sensitive to changes in temperature, pH, and salinity (Ali et al., 2009; Jiao et al., 2010; Upadhyay et al., 2011). Nutrient availability is strongly correlated with microbial EPS production, as the availability of different carbon (C) sources directly influences the precursor molecules available to anabolize into EPS (Celik et al., 2008; Ghosh et al., 2011). Lack of nitrogen (N) or phosphorus (P), inferred from a high C:N ratio, has also been shown to positively affect EPS production of soil microbial isolates (Quelas et al., 2006; Roberson, 1991; Staudt et al., 2012). Thus, microbial production of EPS may be enhanced in marginal soils that inherently possess low N content, but could be limited by C availability given a lack of available organic material.

Cultivation of perennial grasses as cellulosic feedstocks on marginal lands is expected to have a central role in climate change mitigation (Abraha et al., 2019; Robertson et al., 2017). Perennial feedstocks also have neutral C costs, a significant benefit over the C-negative costs of other biofuels such as corn (Gelfand et al., 2011; Tilman et al., 2006). Perennial grasses such as switchgrass (*Panicum virgatum*) possess extensive rooting systems that persist over multiple growing seasons in the soil (Chimento et al., 2016; Ontl et al., 2015, 2013). Roots provide C-inputs to rhizosphere microbial communities in the form of root exudates and mucilage (Mao et al., 2014) and to the total soil community in the form of decomposing root litter (Jackson et al., 1997). These inputs could alleviate C-limitation for microbial communities in marginal soils. Perennial rooting systems can input plant C deeper into the soil profile than do annual plants, potentially increasing soil capacity to sequester C at depth (Anderson-Teixeira et al., 2009; Tilman et al., 2006). Notably, perennial grasses significantly enhance soil aggregation under long-term cultivation (Jastrow et al., 1998; Ontl et al., 2015), and aggregated soils store C more effectively than do soils lacking structure (Liao et al., 2006; McGowan et al., 2019). In addition, studies show that switchgrass (SG) biomass in the field is often not enhanced by nutrient amendments in marginal soils (Brejda, 2000; Parrish and Fike, 2005; Ruan et al., 2016), which makes its cultivation on such soils more cost-effective. This raises the question of whether there are mutualistic relationships between SG roots and their associated soil microbial community that contribute to making up the nutrient deficits in these soils (Rodrigues et al., 2017). In particular, SG may directly facilitate microbial production of EPS by providing microbes with labile C precursors (Mao et al., 2014) and indirectly enhance EPS production by altering soil water potentials through root uptake and potentially through hydraulic lift (Caldwell et al., 1998).

In this study we tested the hypothesis that soil microbial communities in a marginal soil produce EPS in response to limited nutrient and moisture availability. We also asked whether altered EPS production changes soil characteristics, specifically aggregate stability. We used a sandy loam soil depleted in total C, N, and P from a field site in central Oklahoma, where SG is endemic. By manipulating watering regimes and amendments of N and/or P, we assessed how nutrient availability and water stress influenced plant root biomass, EPS, and soil characteristics. We ^13^CO_2_-labeled the switchgrass plants for 12 days to track plant photosynthate C into EPS and bulk soil. Monosaccharide content was analyzed to determine the composition and origin of EPS. Our objective was to determine if SG cultivation can alter microbial activity to enhance beneficial soil characteristics, such as aggregate stability, that are lacking in marginal soils.

## 2. METHODS

### Soil collection and preparation

Soils were collected from a pasture soil in Caddo County, OK, near the town of Anadarko (35.072417/-98.303667). The soil, described by the USDA soil series as Pond Creek fine sandy loam with 1-3 percent slopes, is classified as a superactive, thermic Pachic Argiustoll (Moffatt, 1973). We consider it to be a marginal soil because of its low C content (< 0.4% total C), low nutrient content (< 0.04% total N, < 6 ppm total P), and high (> 70%) sand content in all three observed horizons down to 1 m in depth (**Table S2**). In November of 2016, a backhoe was used to excavate 1-m deep soil pits. In this range, the soil profile was characterized as having three distinct horizons– an A horizon with noticeably more organic material in the top 25 cm, a B horizon with greater sand content from 25-70 cm, and a deeper, noticeably denser C horizon from 70 cm and below. Bulk density cores were also taken from each horizon (**Table S2**).

After removing living plant material from the surface (0-3 cm), approximately ~4 m^3^ of soil was collected from each horizon using a backhoe, and transported to our greenhouse facility at the University of California, Berkeley (UCB). Soils were stored indoors and in December 2016, soil from each horizon was homogenized in a 0.255 m^3^ cement mixer and then stored in sandbags for two months at ambient conditions (25 °C during the day and 18 °C overnight). Sub-samples were taken from each homogenized horizon for initial soil chemistry assays (**Table S2**). Bulk density cores were weighed fresh and dried at 70° C until no change in soil weight was observed to assess field bulk density of each horizon.

### Mesocosm preparation

Based on field observations, soil profiles were re-created with homogenized soil horizons in 30 clear, impact-resistant polycarbonate cylinders (hereafter referred to as “mesocosms”), 122 cm in length and 19.7 cm in diameter. In each mesocosm, we aliquoted the A, B, and C soil to allow each horizon 33 cm of vertical depth, packed at field bulk density. At the base, each cylinder was sealed with a custom-fitted polycarbonate cap and 500 g of coarsegrained sand to provide drainage. During packing, an anion exchange resin membrane (Membranes International, Ringwood, NJ) was added to the center of each horizon to provide an integrated measure of available PO_4_ in that horizon over the course of the study.

To establish five experimental treatments, before adding the A horizon soil to the mesocosm, it was mixed in a cement mixer for 2 minutes with: no additions (control and low water treatments), added N (N), added P (P), or combined N and P (N/P) amendments (**Fig. 1A**). N was added in the form of 0.13 g kg^−1^ dry soil of ESN Smart Nitrogen slow-release coated urea (44-0-0, Agrium) in accordance with the recommendations provided by Oklahoma State’s Division of Agricultural Sciences and Natural Resources for high biomass SG cultivation (Arnall et al., 2018). P was added in the form of 0.48 g kg^−1^ slow-release rock phosphate (0-3-0, Espoma) to bring the total concentration of plant extractable P up to 20 ppm, which resulted in an amendment consistent with manufacturer recommendations as well as Oklahoma State University’s recommendations for SG cultivation in soils with ~5 ppm of total phosphorus.

**Figure 1.**
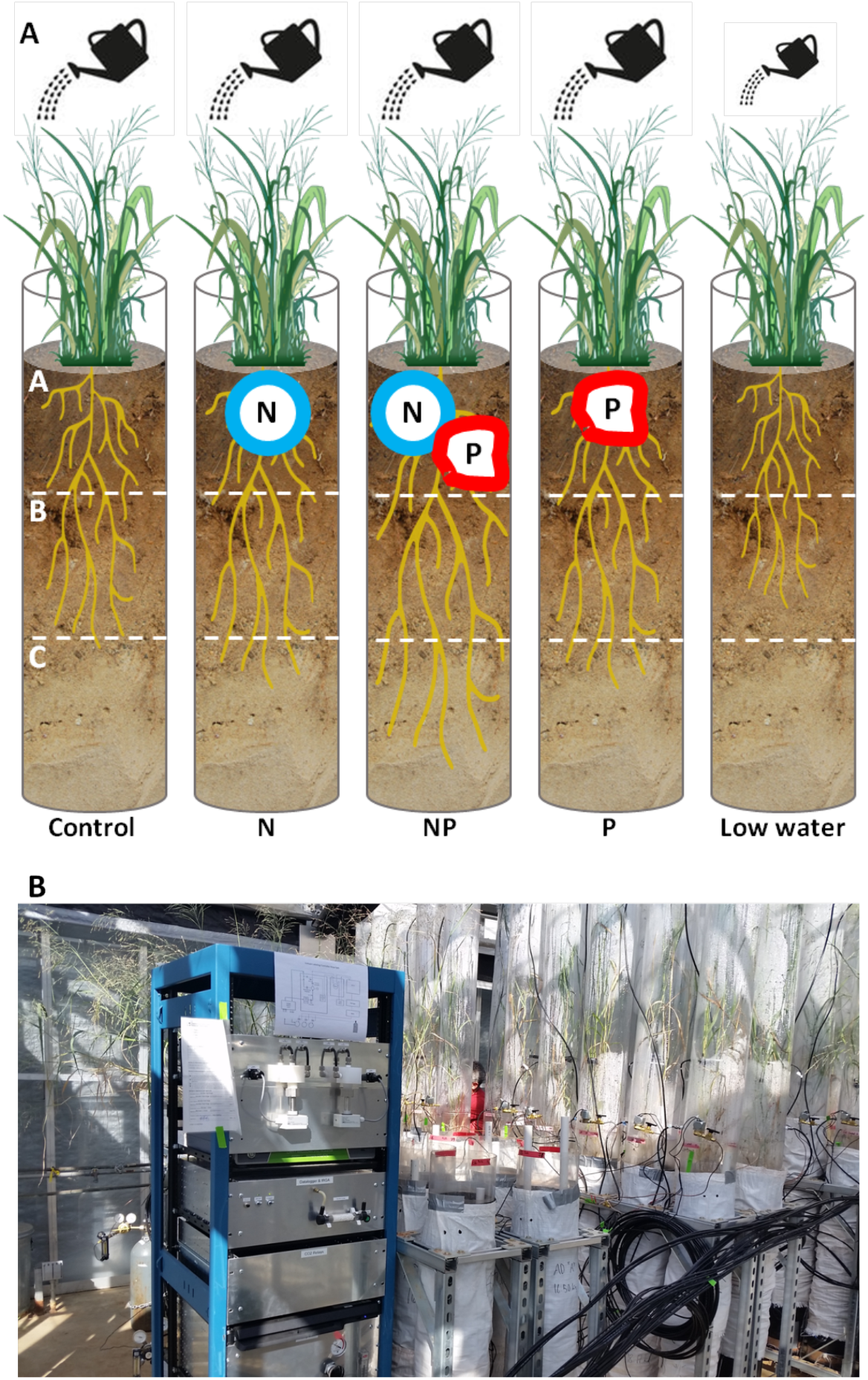
^13^CO_2_ labeling greenhouse experiment designed to test the effects of N and P fertilization and soil water on switchgrass growth, microbial communities and soil properties. A) Schematic of experimental design - switchgrass mesocosms contained three soil horizons (A, B, and C) reconstituted in 1.22 m cylinders, with the A horizon subject to either no treatment (control), nitrogen fertilization by 44-0-0 coated urea (N), phosphorus fertilization by 0-3-3 rock phosphate (P), both nitrogen and phosphorus fertilization (NP) or a 50% reduced watering regime (Low water). Each treatment had 6 replicates, 3 of which were labeled with ^13^CO_2_ and 3 of which were labeled with ^13^CO_2_, for a total of 30 mesocosms. B) A picture of the greenhouse experimental setup showing mesocosms with planted SG, attached labeling chambers and delivery system, and the ^13^C/^12^C CO_2_ gas flow and concentration control panel.

Completed mesocosms were wrapped in black high-density polyethylene sheeting and then white polypropylene sacking (to prevent soil temperatures from being elevated by solar radiation) and stored for an additional five weeks before planting with SG. Soil moisture probes (EC-20; METER Group, Pullman, WA) were installed in the A horizon of the control and low water treatments mesocosms to maintain target moisture conditions. During this period, 2 L of deionized H_2_O was added to each mesocosm every week to re-hydrate the soil profile and allow it to equilibrate before planting. A SG genotype, NFSG 18-01, from the Nested Association Mapping population (NAM) with established high biomass productivity in both Oklahoma and Tennessee was selected for this experiment. SG is highly heterogeneous, and every plant is genetically distinct. To avoid any genetic variation among treatments and replicates, we used a single clone for this experiment. The plant was grown in the Noble Research Institute (TRI) greenhouse at 32°C (daytime) / 21°C (nighttime) and 16 h photoperiod for maximum growth. A total of 120 clonal ramets were prepared from one plant and shipped to UCB. Uniform ramets of NFSG 18-01 were planted into each of the 30 mesocosms in early May 2017. Mesocosms were arranged in 6 wheeled stainless-steel caddies insets of five (one of each treatment, in random order), making each caddy equivalent to a “plot.” Thereafter, mesocosms were watered with 100 ml of deionized H_2_O daily—roughly equivalent to the rainfall experienced in TRI field plots in southern Oklahoma in the higher precipitation months of May and June. After plants were established within the mesocosms (four weeks), watering for the low water treatment was reduced to 50 ml of H_2_O daily. After eight weeks, the temperature in the green house was increased to 32 °C (daytime) / 21 °C (nighttime) to simulate growing season conditions in Oklahoma.

### ^13^CO_2_ pulse-chase labeling

After plants had grown for 18 weeks, we performed a 12-day ^13^CO_2_ pulse-chase labeling. Half (15) of the mesocosms were labeled with ^12^CO_2_ (Praxair, Danbury, CT) as controls for future stable-isotope probing (SIP) and the other half were labeled with 99 atom-percent ^13^CO_2_ (Cambridge Isotope Laboratories, Tewksbury, MA), providing three replicates of each treatment under each labeling regime. Labeling was carried out using a custom apparatus consisting of a Picarro G2131-I Analyzer (Santa Clara, CA) and Infrared Gas Analyzer, (IRGA, Campbell Scientific, Logan, UT) combined with a CR1000 Datalogger (Campbell Scientific, Logan, UT) to enable real-time assessment of [^12^CO_2_] and [^13^CO_2_] in up to 32 chambers (16 of each type of CO_2_).

### Harvest of plant biomass and processing of soils for analysis

At the completion of the pulse-labeling in September, ^13^CO_2_ enriched mesocosms were destructively harvested by clipping SG shoots at the soil surface and partitioning soil horizons for sample collection. For each individual soil horizon, roots were collected from bulk soil by hand, washed in deionized water, and dried at 70° C until no change in root weight was observed. Fresh bulk soil was aliquoted for pH, soil chemistry, and EPS extractions and stored at 4° C. Fresh bulk soil was also aliquoted for assessment of water-stable aggregates and stored in open bags in the greenhouse for air-drying. Gravimetric soil water content was measured for each horizon of each replicate by drying fresh bulk soil at 70°C until no change in soil weight was observed. Soil water content was converted to soil water potential with water retention curves that were generated from samples from each horizon using a pressure plate apparatus (WP4C, METER Environment, Pullman, WA) and a van Genuchten model to apply a non-linear fit to the data (Seki, 2007). Volumetric soil water content measured by EC-20 probes in the A horizon of the control and low water treatments was also converted to soil water potential using these water retention curves.

5 g soil was extracted with 20 ml 0.5M K_2_SO_4_ to assess dissolved organic carbon (DOC) and total dissolved nitrogen (DN). DOC in the extract was measured using a Shimadzu (Kyoto, Japan) TOC-L series CSH/H-type TIC/TOC analyzer as previously described elsewhere (Jenkins et al., 2017). DN in K_2_SO_4_ extracts was measured after combustion by detecting NO with a chemiluminescence gas analyzer. Soil pH values were determined in slurries made by mixing 5g of soil with 5ml of 0.01M CaCl_2_. 1 g dried soil was sent to Oregon State University’s Central Analytical Laboratory (Corvallis, OR) for total C quantification by combustion at 1150 °C on an Elemental Macro Cube. PO_4_ accumulation on ion exchange resin membranes was assessed by extracting anions with 0.5 M HCl and assessing the ppm of P in the extract with a microplate reader using methods described by D’Angelo et al., 2001.

### EPS extraction and analysis

We used a modified cation exchange resin (CER) extraction method (Redmile-Gordon et al., 2014; Wang et al., 2019) to examine how EPS content in bulk soil differed between depths and treatments at the end of the greenhouse experiment The CER reduces binding between multivalent cations and polymeric substances (Sheng et al., 2010), releasing EPS into the extraction buffer solution. This approach minimizes microbial cell lysis that can potentially bias the results (Redmile-Gordon et al., 2014; Steinberger and Holden, 2004; Wang et al., 2019; Zhang and Yan, 2012), and maximizes extracted EPS yield (Frølund et al., 1996; Sheng et al., 2010). Combining this CER extraction with an ethanol precipitation step isolates high molecular weight carbohydrates (Chang et al., 2007), thus targeting soil carbohydrates that are both extracellular and polymeric - i.e., EPS.

In the modified extraction method, the extraction buffer was phosphate buffer saline (PBS; Gibco, Grand Island, NY). Specifically, 5 g soil (kept at 4°C until extraction) and 10 ml phosphate-buffered saline (PBS) were added to a 50 ml tube containing 1 g CER (Dowex^®^ Marathon^®^ C, 20–50 mesh, Na^+^ form, Sigma-Aldrich, St. Louis, MO). This slurry was shaken for 30 minutes at 4 °C and subsequently centrifuged at 3000 rpm for 10 minutes at 4 °C. The supernatant was passed through a 0.2 μm nylon filter, and polysaccharides were precipitated from the filtrate with three volumes of 100% ethanol and concentrated 10x.

To extract EPS for 13C analysis, the extraction procedure was upscaled 18x to obtain enough C in the extract to allow IRMS analysis Extracts were precipitated twice in ethanol, to reduce sample volume to 1 ml. Reduced volume extracts were transferred to small tin cups (Costech, Valencia, CA) and evaporated to complete dryness at 70°C. These tin cups were then folded into tight tin balls for IRMS analysis.

Total EPS was quantified by measuring carbohydrates with a sulfuric acid/phenol method (DuBois et al., 1956), modified for microplates. A colorimetric reaction mix composed of 50 μl of each EPS sample (or standard), 150 μl sulfuric acid (95-98%, A300-212, Fisher Chemical), and 30 μl 5% phenol (Spectrum chemicals) were added to a 1 ml well in a 96 well polypropylene deep-well plate (Thermo Scientific Nunc, Waltham, MA, USA). Plates were tightly covered with polypropylene lid and placed on micro-plate block heater for 45 minutes at 100°C, then allowed to cool for 15 minutes. 100 ml of the mix was transferred from each well of the polypropylene plate to a clear, flat bottom, polystyrene 96 well microplate (Greiner Bio-One) and placed in Spectramax plus 384 plate reader (Molecular Devices) to measure absorbance at 490 nm. Carbohydrate content was measured against a calibration curve of glucose in the range of 0.5-250 μg ml^−1^.

To assess the monosaccharide composition of EPS, 20 μg ml^−1^ solutions of each EPS extract were generated. These solutions were hydrolyzed by adding an equal volume of 4M Trifluoroacetic acid (TFA; Sigma-Aldrich) to attain a 2M final concentration, before being incubated for 90 minutes at 121°C. Hydrolysates were washed twice with isopropanol by evaporating isopropanol with a TECHNE sample concentrator (Cole-Parmer Ltd., UK), and were then eluted with 0.5 ml ultrapure water. Subsequently, re-suspended samples were centrifuged for 10 minutes at 13,000 x g at 4°C, to remove solids, and 80% of the supernatant was collected for analysis. Monosaccharide composition was measured with a Dionex ICS-3000 ion chromatography system with CarboPac™ PA20 column (Thermo Fisher Scientific). Samples were analyzed in two runs with two KOH eluent concentrations, 2mM and 18mM, as at 2mM arabinose and rhamnose peaks overlap while at 18mM xylose and mannose peaks overlap (Yeats et al., 2016). We used this data to calculate the ratio of hexose to pentose sugars in extracted EPS to verify its microbial origin (Gunina and Kuzyakov, 2015; Oades, 1984). Given that it is recommended to assess EPS monosaccharide composition in the context of the plant being studied (Gunina and Kuzyakov, 2015), we also sampled SG roots from TRI’s Red River field site (Burneyville, OK, 33.882235/-97.275919) in early May 2017 to calculate the hexose to pentose ratio in polysaccharides from SG root mucilage. EPS extraction, quantification and monosaccharide composition was done both on roots and bulk soil samples as described above.

### Soil aggregate stability

Soil aggregate stability was measured with a wet sieving method (Kemper and Rosenau, 1986) on 1-2mm soil aggregates that had been sieved from air-dried soil. 5g of these aggregates were placed on 0.25mm mesh sieve and repeatedly dunked in a water cup for 5 minutes, using a mechanical dunking apparatus (Singer et al., 1992). The mass of unstable aggregates (those that dispersed) versus stable aggregates (stayed on the sieve) was dried and measured. The following ratio was used as the measure for soil aggregate stability:

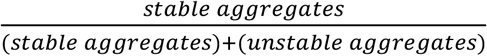

Before assessing soil aggregate water-stability, all soil samples from all treatments were air-dried and had consistent low soil water content to avoid confounding the assay.

### Phospholipid fatty acid analysis (PLFA)

Microbial biomass in each soil sample was determined by Phospholipid Fatty Acid (PLFA) analysis, with a high throughput 96 well plate method to extract and trans-esterify PLFAs, as described by (Buyer and Sasser, 2012). PLFA’s were extracted from 2 g dry soil samples from the greenhouse experiment, in the B and C horizons extracts of two 2 g dry soil samples were combined. After transesterification steps Fatty Acid Methyl Esters (FAMEs) were then analyzed by gas chromatography (Agilent Technologies, Wilmington, DE, USA) and identified using the MIDI Sherlock Microbial Identification System (MIDI, Newark, DE, USA). An internal standard,19:0 phosphatidylcholine (Avanti Polar Lipids, Alabaster, AL, USA) and chromatogram peaks of a PLFAD1 calibration mix and peak library (MIDI Inc, Newark, DE, USA) were used to calculate concentration of analyzed PLFA’s and total microbial biomass.

### ^13^C analysis

^13^C in both bulk soil samples and EPS extracts from the surface horizon were analyzed with an IsoPrime 100 continuous flow isotope ratio mass spectrometer (IRMS) interfaced with a trace gas analyzer (Isoprime Ltd, Cheadle Hulme, UK). ^13^C enrichments were calculated by subtracting ^13^C atom% natural abundance (found in control ^12^C treatments) from the total ^13^C atom% found in ^13^C treatments. ^13^C atom% was multiplied by either total C per g soil for each soil sample, to calculating the amount of labeled soil C, or by EPS per g soil, to calculate the amount of labeled EPS.

### Oklahoma field sampling

We conducted soil coring campaigns to compare EPS content along bulk soil depth profiles of long-term perennial deep-rooting SG fields (10-20 years) and adjacent annual shallow-rooting crop fields. Soil coring campaigns were conducted at TRI’s Red River field site (10-year SG cultivation) and at a field site near Stillwater, OK (20-year SG cultivation, 36.133378/-97.104284). Soil cores were excavated using a Giddings probe (Giddings Machine Company, Windsor, CO). Soil core tubes, ~10 cm diameter and ~1.2 m length (with a 9 cm diameter Zero Contamination system liner), were used to take out soil cores from up to ~3 m depth, for three replicate 1.2 m cores. Each 1.2 m core was cut into three sections of ~30 cm, and soil from the bottom 20 cm of each section was stored at 4°C for one week before EPS was extracted and quantified (as described above).

### Statistical analysis

Statistical analysis and data visualization were conducted with R version 3.6.0 (R Core Team, 2018). Significant differences in soil properties and SG root biomass between treatments and soil horizons for the greenhouse study and differences in EPS content between SG and annual crop fields with depth were determined by ANOVA. Every two adjacent depths from the field samplings were combined during analysis to increase statistical power. Significant differences in microbial biomass between treatments and soil horizons were assessed for the A and B horizon only, given the prevalence of N/A results for PLFA microbial biomass in the C horizon. For further multiple comparisons between treatments and soil horizons, pairwise t-tests were conducted without pooled standard deviations (Welch’s t-test), as the assumption of equal variance between samples for Tukey’s test were not met for some of the analyses. To correct for multiple testing, the Benjamini-Hochberg correction was used (Benjamini and Hochberg, 1995). Box-whisker graphs were built with the ggplot2 package (Wickham, 2016) in R. Significant differences in EPS monosaccharide composition as a result of treatment were determined by MANOVA.

To determine which measured properties best explain EPS and soil aggregate stability variability between treatments and horizons, we employed multiple linear regression analysis. Multicollinearity between examined factors was detected by calculating the variance inflation factor (VIF) between them (Fox and Weisberg, 2011), and removing highly correlated factors with VIF value above 3 (Zuur et al., 2010) in a stepwise manner. After removing highly correlated factors, factors with low explanatory significance to the multiple linear regression models were removed after Akaike information criterion (AIC) estimation of the relative quality of the models (Venables and Ripley, 2002). Correlation between factors was visualized with correlation matrix charts (Peterson et al., 2018) (presented in the Supplemental Online).

We used path analysis, conducted with the lavaan package (Rosseel, 2012) in R, to assess how root biomass and other measured soil factors interact to affect observed EPS content and the percentage of water-stable aggregates. Although path analysis is built for larger sample sizes than we have in our study, it provides conservative fit estimates and is not prone to Type 1 errors (Shipley, 2016). Using a workflow based on that presented in (Petersen et al., 2012), we developed a full model of interacting paths between root biomass, soil water potential, pH, microbial biomass, DOC, DN, PO_4_ accumulation, EPS content, and the percentage of water-stable aggregates based on theoretical linkages between the relevant measured variables (**Figure S4**). We iteratively removed non-significant edges between the measured soil factors (p < 0.1) from the resulting path model until all edges were significant, and evaluated the fit of this reduced model to the data using a X^2^ test and the Tucker-Lewis index. Visualization of the resulting path analysis was performed using the *semPlot* package (Epskamp, 2019) in R.

## 3. RESULTS

### EPS and soil factors

EPS concentrations in mesocosm soils after 143 days of SG growth varied significantly as a function of both treatment (ANOVA, F = 5.16, D.F. = 4, P = 0.001) and depth (ANOVA, F = 238, D.F. = 2, P < 0.001; **Fig. 2A**). In all treatments, EPS content was greatest in the surface A horizon; ranging from 11.74 ± 2.04 μg g^−1^ (mean ± SD) in the A horizon to 6.99 ± 1.60 μg g^−1^ and 2.28 ± 1.43 μg g^−1^ in the B and C horizons, respectively (**Fig. 2A**). EPS content only differed between treatments in the A horizon (ANOVA, F = 8.24, D.F. = 4, P < 0.001; **Fig. 2A**), and these differences were primarily driven by the enhanced EPS content observed in the NP treatment relative to all other treatments (Welch’s t-tests, P < 0.025).

**Figure 2.**
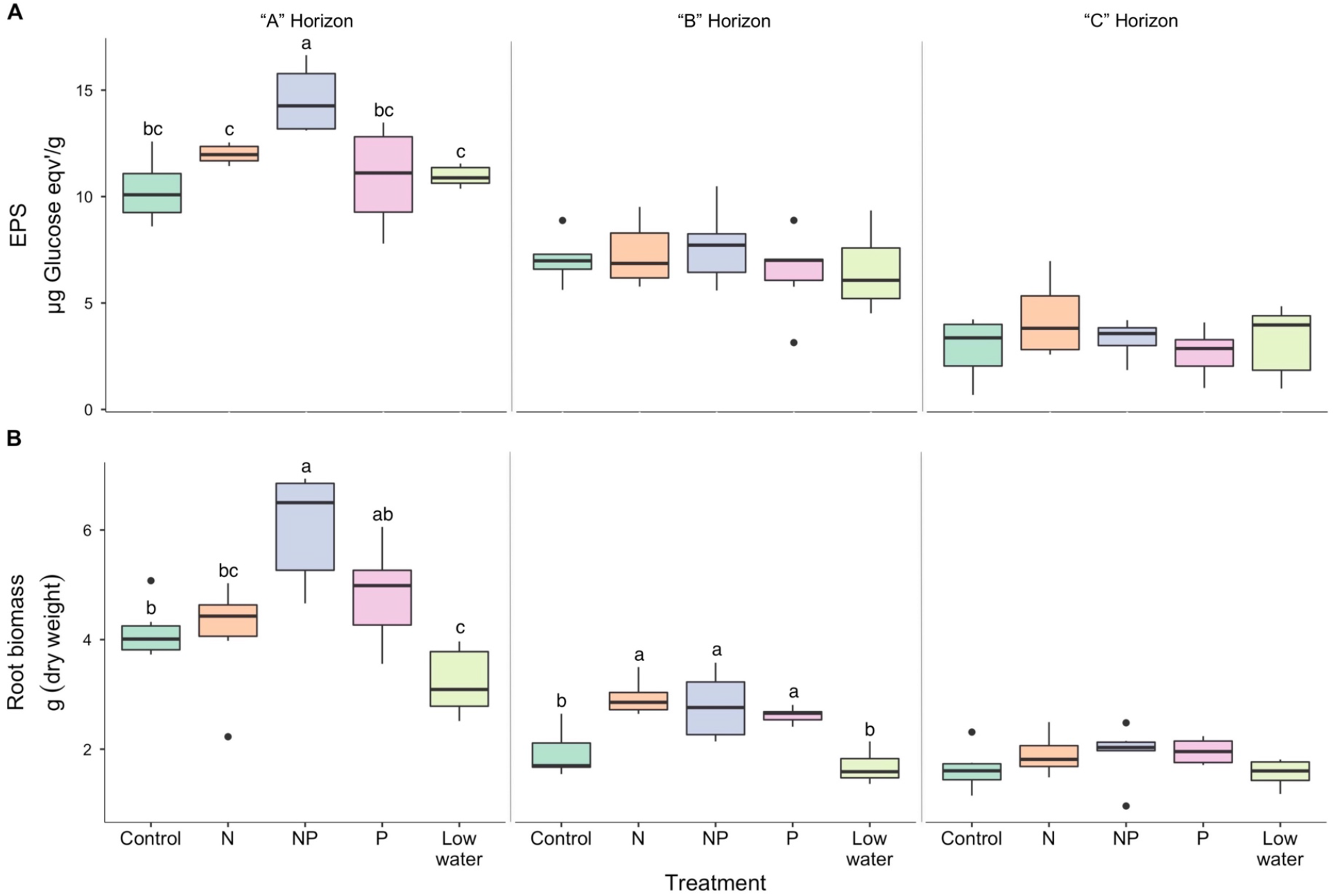
EPS and root biomass in switchgrass (SG) microcosms with five fertilizer/moisture treatments and three soil horizons after 140 days of growth. Box-whisker plots of A) EPS content (glucose equivalent, micrograms per gram dry soil) and B) SG root biomass (grams dry weight) recovered from bulk soil by treatment and horizon. Lowercase letters indicate significant differences between treatments within a horizon (Welch’s t-test P < 0.05, after Benjamini-Hochberg correction for multiple testing). n = 6 per treatment/horizon.

Root biomass exhibited similar trends as EPS - the greatest root biomass was observed in the A horizon, with reduced biomass in the B and C horizons (**Fig. 2B**). Root biomass varied significantly by treatment in the A horizon (ANOVA, F = 9.64, D.F. = 4, P < 0.001; **Fig. 2B**), and was highest in the NP treatment. Root biomass also varied significantly by treatment in the B horizon (ANOVA, F = 12.36, D.F. = 4, P < 0.001; **Fig. 2B**), mostly as a result of higher root biomass observed in fertilized treatments.

DOC exhibited similar trends to both EPS and root biomass: DOC was highest and varied significantly between treatments in the A horizon; it was significantly highest in the N and NP treatments relative to all the others (Welch’s t-tests, P < 0.019) (**Fig. S1A**). DN was significantly higher in the A and B horizon of both nitrogen-amended treatments. DN was also significantly higher in the N treatment relative to the NP treatment, possibly indicating greater demand for N in the higher root biomass NP treatment (**Fig. S1B**). Accordingly, N fertilized treatments had the lowest ratios of DOC to DN (dissolved C/N, **Fig. S1C**). In the A horizon, the N and NP treatments had significantly lower pH than non-N treatments (**Fig. S1D**). In all three soil horizons, soil water potential (measured at the time of the destructive harvest) was significantly lower in both N-fertilized treatments and the low-watering regime treatment relative to the control and P treatments (**Fig. S1E**). PLFA-measured microbial biomass was significantly affected by soil depth, with minimal biomass in the B horizon and barely detectable biomass in the C horizon (F = 349, D.F. = 1, P < 0.001). We did not observe significant treatment effects on microbial biomass across horizons (F = 2.43, D.F. = 4, P = 0.065; **Fig. S1F**) or within the surface horizon (F = 1.03, D.F. = 4, P = 0.414).

We compared the influence of measured soil properties on EPS across all treatments and horizons using multiple linear regression. Many factors were significantly correlated with one another (**Fig. S2**, **S3**); the most collinear factors were removed from the analysis according to their VIF (as explained in the Methods section). The resulting model explains a large proportion of the variation in EPS (R^2^ = 0.799) between treatments and horizons; root biomass, soil water potential, DN and microbial biomass were the most significant explanatory factors (P = 0.010, 0.034, 0.010 and < 0.001, respectively; **Table 1**). Because our dependent variable, EPS, only varied significantly between treatments in the A horizon, we did not investigate interactions between treatment and depth in other variables, and we employed a second model for only the A horizon. This model also succeeded in capturing a majority of the variability in observed EPS between treatments (R^2^ = 0.667); root biomass was again the most significant factor (P < 0.001), and soil water potential and pH were additional significant factors (P = 0.007 and 0.009, respectively). A final model was developed with N addition included as a confounding factor to account for decreases in pH as a result of N fertilization (**Fig. S1D**). In this model root biomass and soil water potential were the most significant factors controlling EPS (P < 0.001 and P = 0.017, respectively), and pH was no longer significant (P = 0.198).

**Table 1:**
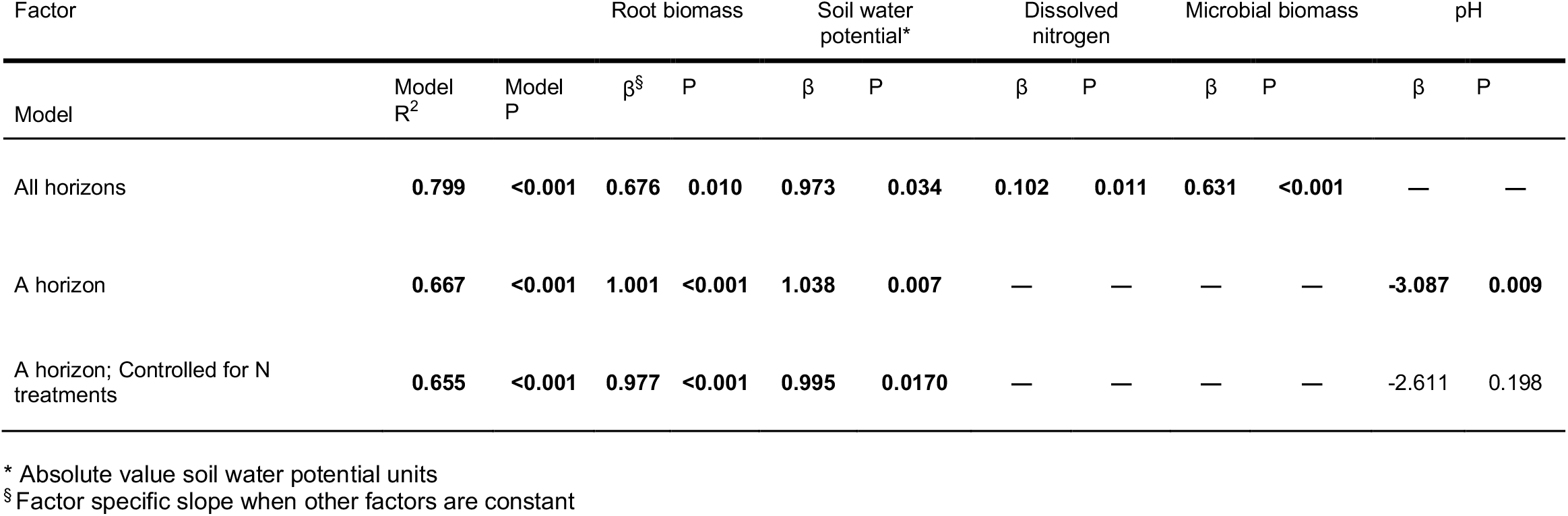
Multiple linear regression models describing relations between most highly explanatory soil factors and EPS, after removing collinear explanatory soil factors with variance inflation factor values above 3 (Zuur et al., 2010).

### ^13^C-labeled EPS and total soil carbon

To assess the proportion of total soil C that was EPS, we expressed soil EPS content as a fraction of total soil C. We found ~0.3% of soil carbon is EPS, with no significant differences between treatments and horizons. Dividing the ^13^C-labeled C observed in EPS by the total ^13^C-labeled C found in the bulk soil (**Table S2**) revealed that 0.18% of newly-fixed plant-derived C in soil was or had been assimilated into EPS. There were no significant differences in the proportion of freshly fixed C recovered in EPS between treatments. We obtained this data only for the A horizon, as a substantial amount of soil was needed to extract sufficient EPS for ^13^C IRMS analysis and significant differences in EPS content were not found between treatments in the B and C horizon.

### EPS monosaccharide composition

We analyzed the monosaccharide composition of soil EPS in the A horizon to assess its potential origin by calculating the ratios of recovered galactose + mannose (G+M, microbially derived) to arabinose + xylose (A+X, plant derived). We found this ratio to be consistently above the accepted cutoff of 2.00 in our samples (average of 3.92 ± 0.25), indicating EPS had a likely microbial origin (Chenu, 1995; Gunina and Kuzyakov, 2015; Oades, 1984). The observed ratio was significantly lower in the A horizon of the NP, N and low-watering regime treatments relative to the control and P treatments ((3.78 ± 0.19, mean ± SD across treatments) vs. (4.14 ± 0.15), respectively; P < 0.02) (**Table S3**). Given that SG root mucilage itself appears to have high galactose content resulting in a (G+M)/(A+X) ratio of 1.45 ± 0.17, we also employed a more conservative M/(A+X) ratio to confirm that the majority of the EPS we extracted was most likely microbial in origin. Recovered EPS still had a value above 2.00 with this modified ratio (2.24 ± 0.13, on average), giving us confidence in our prior conclusion. This more conservative ratio did not differ between treatments in the A horizon. Monosaccharide composition was found to vary significantly as a result of treatment (MANOVA F = 6.34, D.F. = 20, P < 0.001).

### Soil aggregate stability and relationship to measured variables

We measured the percentage of aggregates that were water-stable to assess the effects of treatment and EPS content on soil aggregation. The percentage of water-stable aggregates was significantly higher in the NP treatment (**Fig. 3**), and we measured a significant positive correlation between soil EPS and the percentage of water stable aggregates (Pearson R = 0.44, P = 0.017; **Fig. S3**).

**Figure 3.**
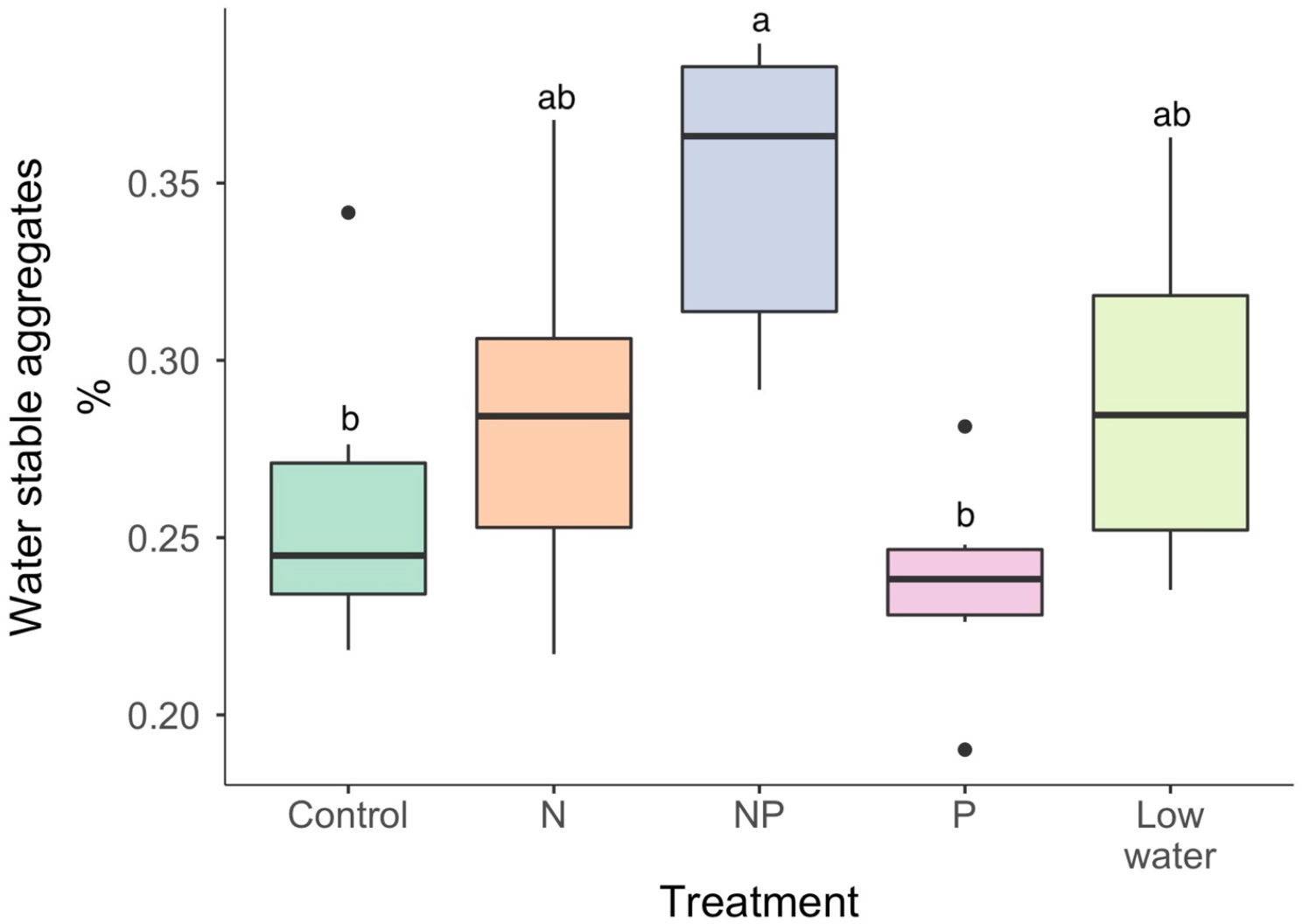
Soil aggregate stability in the surface horizon of switchgrass (SG) mesocosms with five fertilizer/moisture treatments after 140 days of growth. Box-whisker plot of the percentage of aggregates recovered from bulk soil that were water-stable in the A horizon. Lowercase letters indicate significant differences between treatments (Welch’s t-test P < 0.05, after Benjamini-Hochberg correction for multiple testing). n = 6 per treatment.

We conducted path analysis to determine how root biomass and our observed soil characteristics may interact to impact both EPS content and the percentage of water-stable aggregates in our mesocosms. Our full model (including all of the soil variables measured) fit the data well according to the model chi-squared statistic (**X**^2^ = 5.474, D.F. = 4, P = 0.242), as did the reduced model where we removed non-significant edges (**X**^2^ = 14.066, D.F. = 21, P = 0.867). The Tucker-Lewis index, which is more sensitive to the number of parameters included in the analysis, indicated that our reduced model fit the data very well (TLI = 1.105, above the 0.9 threshold). Given that the reduced model is a nested variant of the full model, we verified that the reduced model did not fit the data in a significantly different manner from the full model using a maximum likelihood ratio test (**X**^2^ difference = 8.592, D.F. = 17, P = 0.952). The reduced model shows that root biomass affects soil EPS content both directly and through the DOC pool, whereas soil water potential acts separately on both EPS and the percentage of water-stable aggregates (**Fig. 4**). Soil aggregation is also affected by pH. In addition, EPS and water-stable aggregates co-vary positively with one another.

**Figure 4.**
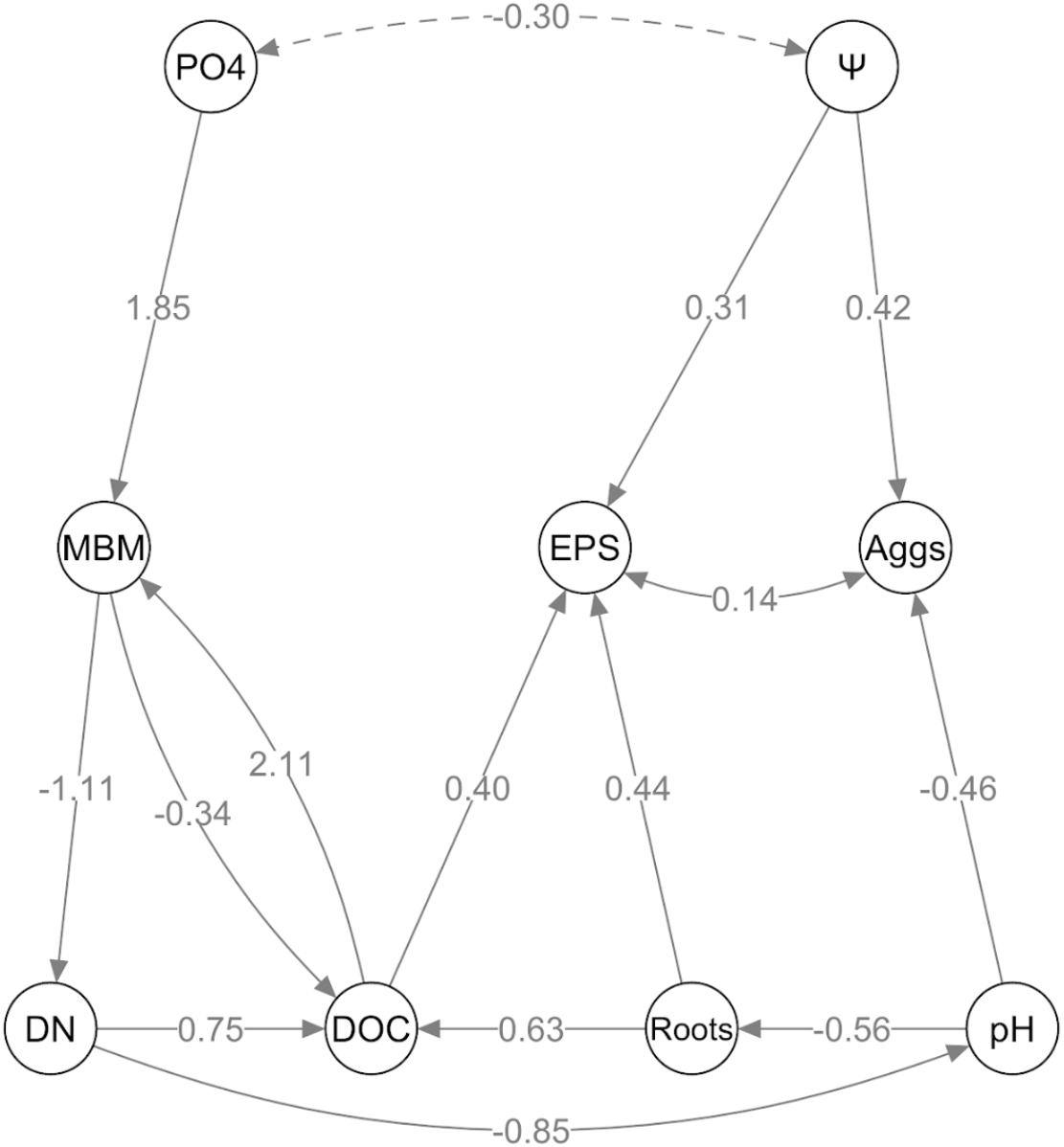
Path analysis of soil factors affecting EPS and soil aggregate stability in the surface soil horizon of SG mesocosms with five fertilizer/moisture treatments after 140 days of growth in a ‘reduced model’, where only significant edges (P < 0.05) are retained. Node labels correspond to the following measured variables: EPS content (EPS), frequency of water-stable aggregates (Aggs), soil water potential (Ψ), pH, SG root biomass (Roots), dissolved organic carbon (DOC), total dissolved nitrogen (DN), microbial biomass measured by PLFA (MBM), and phosphate accumulation on anion exchange membranes over the course of the study (PO_4_).

### EPS in soil core field sampling

To assess the impact of deep-rooted perennial grass cultivation on field stocks of soil EPS, replicate soil cores were collected from 10- and 20-year-old SG marginal soil fields in Oklahoma and compared with cores from adjacent fields that had been historically managed with rye (Red River) or wheat and sorghum (Stillwater) row crops under consistent tillage. Significantly larger stocks of EPS were observed in the SG field soils compared to those observed under annual crops (Red River F = 11.58, D.F. = 4, P < 0.001; Stillwater F = 14.88, D.F. = 4, P < 0.001; **Fig. 5**). This significant enhancement of EPS content extended over 1.5 m deep in the soil, with concentrations of ~10 μg g^−1^ in the surface layers and ~2 μg g^−1^ below 180 cm depth soil samples (Red River F = 2.46, D.F. = 4, P = 0.062; Stillwater F = 4.43, D.F. = 4, P = 0.005; **Fig. 5**).

**Figure 5.**
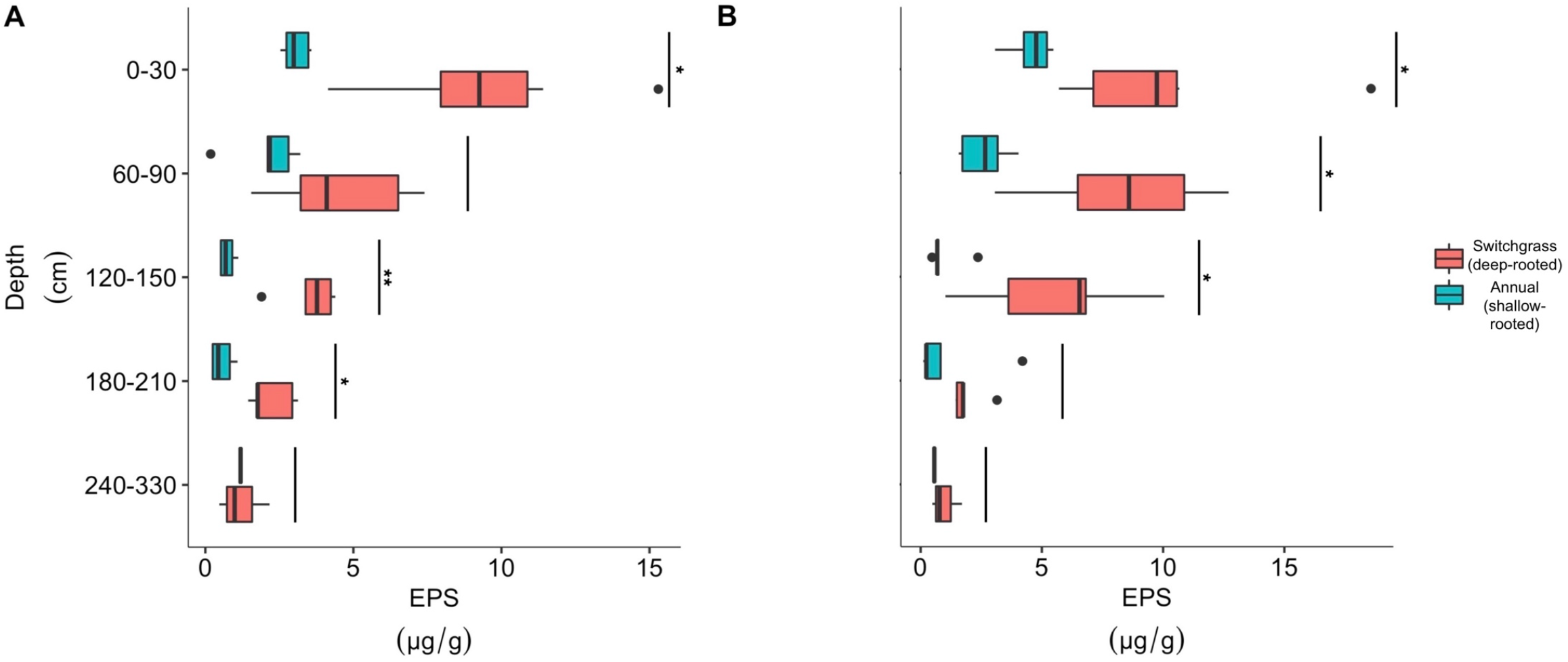
Soil depth profile of EPS content in switchgrass (SG) and annual crop fields. Box-whisker plots of EPS content (glucose equivalent micrograms per gram dry soil) in deep soil cores from plots subjected to A) 20-year (Stillwater, OK) and B) 10-year (Red River site, OK) no-till cultivation of deep-rooted SG compared to paired plots at each site planted with short-rooted annual rye or wheat/sorghum and managed with annual tillage. Asterisks indicate significant differences between deep-rooted Switchgrass plots compared to paired annual plots at each sampled depth (Welch’s t-test, P < 0.05 for * and < 0.01 for **, after Benjamini-Hochberg correction for multiple testing). n = 6.

## 4. DISCUSSION

### Nutrient and water treatment effects on switchgrass root biomass and EPS

Our results demonstrate that SG root biomass is a major driver of soil EPS content under abiotic stress in our marginal soil. Water stress is also a significant driver of soil EPS content (**Table 1**) and the EPS produced under water stress may improve water-stable aggregation in a marginal soil (**Fig. 3**). However, our results did not support our hypothesis that soil microbes exposed to greater nutrient limitation would produce more EPS. Instead, we found that the most important factor affecting EPS production was SG root biomass, which itself was enhanced by fertilization with N and P (**Fig. 2**). Our observation that dissolved C:N ratios are negatively correlated with EPS (**Fig. S2**, **S3**) contrasts with previous findings that high C:N ratios enhanced EPS production (Pal and Paul, 2013; Roberson, 1991; Sheng et al., 2006, p. 200; Wang and Yu, 2007). However, these previous studies were performed with microbial isolates in culture rather than direct soil observations. Data about microbial EPS production in response to N limitation in soil is scarce, though it has been shown that specific N management practices can increase or decrease the quantity of EPS-like carbohydrates bound to the soil heavy fraction, depending on the quantity of N added (Roberson et al., 1995).

More recently, Redmile-Gordon et al. showed that high C availability increased soil extracellular polysaccharide content (Redmile-Gordon et al., 2015). This suggests that the availability of C precursors for EPS production may be the limiting factor in soil environments, a condition that is likely important in marginal soils with low C content. We have evidence that enhanced root biomass may have provided C precursor compounds to the soil microbiota (Cheng and Gershenson, 2007; Zhalnina et al., 2018), as DOC was enhanced in treatments that had high root biomass and EPS (**Fig. S1A**). Root biomass and DOC also had the highest correlation coefficients of any measured variables with EPS content across all treatments in the A horizon, (Pearson R > 0.68; **Fig. S3**), and were also significantly correlated with one another (Pearson R = 0.51; **Fig. S3**). Our path analysis also provides support for this hypothesis, as root biomass influenced EPS strongly through the DOC pool (**Fig. 4**). We should note that our measurements of DOC and EPS likely overlap to some extent, although our EPS extraction method targets polymeric carbohydrates bound within the soil matrix (Wang et al., 2019) while DOC assays target soluble C compounds.

While several field studies suggest that SG productivity is relatively insensitive to N-fertilization (Brejda, 2000; Pedroso et al., 2011; Ruan et al., 2016; Thomason et al., 2005), SG root biomass clearly responded to the NP treatment in our soil. This is not entirely surprising given the highly N- and P-deplete character of the marginal soil used in this experiment (**Table S2**). The literature suggests there are threshold values of N availability, below which N amendment can enhance SG biomass (McGowan et al., 2018; Vogel et al., 2002). It is also possible that P-limitation may have played a role in the lack of SG biomass response to our N-only amendment; we observed a higher increase in root biomass in our +NP treatment relative to the +N treatment (**Fig. 2A**), and higher levels of dissolved N consumption in the +NP treatment (**Fig. S1B**).

Root biomass may also alter EPS production by reducing soil water potential and increasing its diurnal variability (Caldwell et al., 1998; Kirkham, 2005) such that EPS production may have been promoted by increased water stress (Roberson and Firestone, 1992). Indeed, the next most significant predictors for EPS production in the A horizon were soil water potential and pH (**Table 1**). The role of pH is likely a by-product of our coated urea N amendment, which likely provided ammonia for nitrifying bacteria to oxidize, releasing protons and acidifying the soil in the process (Robertson and Groffman, 2015; Zhu et al., 2016). For this reason, we also employed a multiple linear model to assess EPS variability between A horizon samples that accounts the effects of N amendment (i.e., changes in pH solely due to fertilization). Once “N amendment” is controlled for as a factor, the effect of pH on EPS drops out as a significant predictor, leaving only root biomass and soil water potential (**Table 1**).

The role of EPS in enhancing microbial resistance to low water potential has been extensively discussed in the literature (Costa et al., 2018; Or et al., 2007; Schimel, 2018), with most data derived from studies on isolates (Chang et al., 2007; Ophir and Gutnick, 1994; Roberson and Firestone, 1992). In our study, soil water potential was significantly correlated with EPS content, in a manner consistent with previous research on the effects of water stress on microbial EPS production (Roberson and Firestone, 1992; Sandhya and Ali, 2015). Furthermore, a recent study of EPS accumulation in soil found that EPS production was reduced in plots that received more water (Marchus et al., 2018); when plant cover was removed from wetter plots, less EPS was observed. It is well known that actively evapotranspiring plants with dense root systems can cause daily changes in soil water potential (Caldwell et al., 1998; Kirkham, 2005). Though our soil water potential measurements at harvest were unable to capture these diel dynamics, we did make continuous soil water potential measurements in our control and low water treatments for the 12 days when they were being labeled with ^12^C or ^13^C CO_2_ (and not being watered) (**Fig. S5**). These data show the daily changes in soil water potential that occurred solely because of root uptake and evapotranspiration, and indicate that 1) our mesocosms experienced diurnal variation in soil water potentials in the A horizon, and 2) there was a greater amplitude in this diurnal variation in soils that were drier (**Fig. S5**).

### Quantifying soil EPS and validating its microbial origin

The quantity of EPS we recovered is low compared to some previous studies, reaching maximum values of 14.6 ± 1.6 μg g^−1^ in the A horizon of the NP treatment (**Fig. 2A**). Recovered EPS was significantly lower in the two deeper soil horizons, and did not vary between treatments at those depths, mirroring the lower microbial biomass in the deeper horizons (**Table 1**; **Figure S1F**). In a recent watering manipulation experiment in an annual grassland, significantly higher amounts of EPS were recovered (150-300 μg g^−1^) using a hot-water extraction method (Marchus et al., 2018). Our choice of extraction method is unlikely the cause of our lower values, however, as our observed EPS content is also noticeably reduced relative to the 170-460 μg g^−1^ from another annual grassland extracted using the same CER technique (Redmile-Gordon et al., 2014). Instead, the low microbial biomass (**Fig. S1F**), very low total carbon content (**Table S2**) and the low availability of DOC (**Fig. S1A**) in our marginal soil likely constrained the amount of EPS that could be produced by local microbial communities. After converting to similar microbial biomass units (Bailey et al., 2002), we found noticeably lower (~70 μg C g-^1^) microbial biomass in our soil than did Marchus et al. (~200–300 μg C g^−1^). Notably, that study also found significant correlation between microbial biomass and EPS content (Marchus et al., 2018). Carbohydrate content and microbial biomass vary extensively across soil types; while our soils have low microbial biomass and EPS content, our observed values fall well within the wide range found in a study of 108 arable, grassland, and forest soils (Joergensen et al., 1996).

Given that switchgrass root biomass was the strongest determinant of soil EPS content, we wanted to know whether the EPS being produced was of direct plant origin or derived from soil microorganisms that were utilizing carbon-rich exudates released by roots. We assessed the likely origin of EPS by comparing the ratio of common hexose sugars associated with microbial EPS—galactose and mannose—to common pentose sugars associated with plant mucilage—arabinose and xylose. The value of the galactose + mannose / arabinose + xylose ratio we measured (3.9 ± 0.2, **Table S3**) indicates that most of the polysaccharides within the EPS we extracted were microbial in origin (Oades, 1984). However, a recent meta-analysis suggests that different plant species may vary significantly in the contribution of galactose to their polysaccharide content, and recommends that polysaccharide ratios be developed specifically for the plants employed in a study (Gunina and Kuzyakov, 2015). We assessed SG root mucilage and found that its galactose content is quite high, so we removed galactose from the hexose:pentose ratio for a more conservative index. Our resulting ratio (2.24 ± 0.13) still exceeds the commonly accepted threshold value of 2.00, giving us some confidence that EPS produced over the course of our study is primarily microbial in origin (**Table 1, Fig. 2**).

### Soil aggregate stability controlled by same factors as EPS production

The formation of water stable aggregates is considered a positive indicator for both soil health and C sequestration (Costa et al., 2018; Rinot et al., 2019; Six et al., 2000; Six and Paustian, 2014). The possibility that microbial polysaccharides can increase soil aggregation was suggested as early as the 1940s (Martin, 1946). EPS is hypothesized to contribute to soil aggregation by providing an adhesive to “glue” soil particles together, increasing both the size and stability of soil aggregates (Tisdall and Oades, 1982). Soil aggregate stability has also been previously linked to the presence of SG roots (McGowan et al., 2019; Tiemann and Grandy, 2015), and there is significant evidence that perennial grasses can enhance soil aggregation (Chimento et al., 2016; McGowan et al., 2019; McLauchlan et al., 2006; O’Brien and Jastrow, 2013). Dense root systems such as those under perennial grasslands may enhance the wetting and drying cycles of soil (as they did in our study; **Fig. S5**), which can also enhance aggregate stability depending on the type of clays present (Singer et al., 1992). Our results show that the percentage of soil aggregates that are water-stable is significantly correlated with EPS content (Pearson R = 0.44; P < 0.05), which itself is best predicted by root biomass, but pH and soil water potential were found by path analysis to be the best predictors of aggregate stability (**Table S1**). This suggests that soil conditions which regulate microbial EPS production are also the primary controllers for aggregate stability in our study.

The results from path analysis (**Fig. 4**) support the conclusion that root biomass exerts strong control over EPS content both directly and indirectly (through the DOC pool), while soil water potential exerts direct control over both EPS content and aggregate stability. Furthermore, aggregate stability and EPS content co-vary, a plausible result given that EPS is thought to promote aggregate stability while aggregates themselves can occlude soil organic C (Jastrow et al., 2007; Six et al., 2002, 2000). We found that EPS monosaccharide composition varied between our treatments (F = 6.34, D.F. = 20, P < 0.001), but further studies are required to determine if differences in EPS composition can affect the formation of water-stable aggregates. Soil pH is often overlooked as a factor that can contribute to soil aggregation (Sollins et al., 1996), but it was also a significant controller of aggregate stability according to our path analysis. There is prior evidence from tropical soils that low pH can enhance aggregate stability (Idowu, 2003; Russell et al., 2018), and some studies suggest that increased soil pH enhances clay dispersion and reduces aggregate stability (Amézketa, 1999).

### Carbon flow from plant photosynthate into microbial EPS

Our results provide evidence of microbial incorporation of C into extracellular polysaccharides, but the total amount of EPS in our mesocosms was lower than that found in other studies. Estimates of carbohydrate content in various soils range up to 10% of total C (Cheshire, 1979), but we observed that the EPS found in our marginal soil after incubation with SG formed only 0.3% of the total soil C pool (**Fig. 2A** and **Table S2**). However, because this fraction of the soil C pool was responsive to our treatments and appeared to exert some control over the formation of water-stable aggregates, it highlights the importance of assessing EPS stocks in soils. This may be particularly true in marginal soils, where the overall depletion of organic C in the surrounding soil environment may enhance the effect of a small pool of actively synthesized polysaccharides that can alter soil characteristics and microbial viability.

The ^13^CO_2_-labeling in our study allowed us to determine that after 12 days of plant exposure to ^13^CO_2_, 4.39 ± 3.72 μg g^−1^ of total soil C contained ^13^C, with no significant differences between treatments (n = 3 for each treatment). ^13^C content in EPS per gram of soil reached much lower values of 0.006 ± 0.003 μg g^−1^, with the least enriched EPS observed in the low water treatment and the most enriched EPS observed in the +N treatment (0.003 ± 0.001 and 0.009 ± 0.001 μg g^−1^, respectively). This indicates that alleviating N-limitation may free the plant to provide photosynthate C to the microbial community, whereas under water-stress, SG may reduce C flow to the rhizosphere. The ratio between labeled EPS to the total ^13^C labeled soil C was ~ 0.18%, with no significant differences between treatments. This ratio indicates how much of the recently fixed C exuded from the roots into the soil was incorporated into EPS by soil microorganisms during the 12 days of labeling at the end of the plant growth period. To our knowledge, no other studies have examined the fraction of EPS produced using freshly fixed plant photosynthate, making it difficult to place our results in context. Future experiments taking advantage of isotope-enabled approaches and labeling systems that have emerged during the last decade (Pett-Ridge and Firestone, 2017) will enable us to evaluate the magnitude of this ratio and how this aspect of plant-microbe interaction may vary between plant species, soil types and abiotic stress conditions.

### Higher EPS in SG fields than adjacent annual grass fields

The data collected from our greenhouse study highlight how SG roots may modify soil conditions to enhance microbial EPS production, and establish that SG photosynthate C can be found in EPS recovered from a marginal soil. However, this data represents greenhouse conditions over a limited period of time (< 1 growing season). Several studies have established that soil organic C is generally enhanced under long-term SG cultivation when compared to paired annual crop fields (Chimento et al., 2016; Dou et al., 2013; Ma et al., 2000; McGowan et al., 2019), but we are not aware of other studies assessing EPS stocks under SG. To assess the field-relevance of our greenhouse study, we determined EPS in soils under long-term (>8 years) SG cultivation vs. long-term annual crop cultivation at two marginal soil sites in Oklahoma. Our results clearly indicate that SG cultivation enhances stocks of EPS (**Fig. 5**), and these enhanced EPS stocks extend more than a meter deep within the soil profile. Root density was also significantly enhanced down to at least 30 cm deep under SG compared to annual crops (data not shown). This indicates that SG cultivation can have significant effects on soil C characteristics much deeper than annual plants appear to reach. If these effects are related to those observed in our greenhouse experiment, then increased EPS under SG cultivation in the field could be indicative of (or a driver of) increased soil aggregation. Enhanced aggregation could then provide a mechanism for the persistence of C under SG cultivation (Liao et al., 2006; McGowan et al., 2019). While the overall sustainability of SG cultivation is a function of many agricultural ecosystem characteristics (trace gas production, fertilization and associated N and P loss to water systems, etc.), the long-term impacts on soil C retention and soil structure are important indices of ecosystem sustainability.

## 5. CONCLUSIONS

We found that SG cultivation can enhance microbial EPS production in a marginal soil. SG root biomass enhances the availability of organic C compounds, providing precursors for microbial EPS production. Root biomass and soil water potential combined to exert significant control over microbial EPS production as well as water-stable aggregate formation. Growing roots absorb water from the soil, providing a mechanism for indirect root enhancement of microbial EPS production and water-stable aggregation formation. We also found evidence of significantly enhanced EPS stocks under long-term SG plots in two field plots, suggesting that these mechanisms could be broadly relevant. Our results add to a growing consensus that SG cultivation can significantly enhance soil characteristics that are of great import to proponents of soil health, especially for marginal lands. More research is required to determine how microbial communities under SG process rhizodeposits into EPS and how this EPS translates to beneficial soil characteristics, as well as broader scale field-studies to assess at what rates EPS accumulates under relevant land management practices.

## Acknowledgments

This research is based upon work supported by the U.S. Department of Energy Office of Science, award DE-SC0014079 to UC Berkeley and subcontracts to the Noble Research Institute, the University of Oklahoma and Lawrence-Livermore National Laboratory (award SCW1555). Work at LLNL was conducted under the auspices of DOE Contract DE-AC52-07NA27344. YS was supported by postdoctoral fellowship grant no. 2016-67012-24717 from the USDA National Institute of Food and Agriculture. We thank David Orme for allowing us to collect soil from his ranch in Anadarko Oklahoma, and Hugh Aljoe, Kelly Craven and Shawn Norton for facilitating site access and soil collection. We thank Yanqi Wu for the opportunity to sample Oklahoma State’s field site near Stillwater, OK. The custom ^13^CO_2_ labeling system was designed by Don Herman. Thanks also to Katerina Estera-Molina, Anne Kakouridis, Sarah Baker, Steve Blazewicz, Evan Starr, Ka Ki Law, Mengting Yuan, Heejung Cho, Alexa Nicholas, Eoin Brodie, Peter Nico, Caleb Herman, Ashley Campbell, and Amrita Bhattacharyya for their help with the destructive harvest of mesocosms; Yuan Wang, Na Ding, Travis Simmons, Josh Barbour, Mellisa McMahon, Jialiang Kuang, Colin Bates, Ryan Gini and Noah Sokol for their help with deep core soil sampling; Madeline Moore, David Sanchez, Cynthia-Jeanette Mancilla and Ilexis Jacoby for their help with processing and soil chemical analyses; and Trent Northen and Kate Zhalnina for their help with TOC/TN analysis.

## REFERENCES

Abraha, M., Gelfand, I., Hamilton, S.K., Chen, J., Robertson, G.P., 2019. Carbon debt of field-scale conservation reserve program grasslands converted to annual and perennial bioenergy crops. Environmental Research Letters 14, 024019. doi:10.1088/1748-9326/aafc10

Adessi, A., Cruz, R., Carvalho, D., Philippis, R.D., Branquinho, C., Marques da Silva, J., 2018. Microbial extracellular polymeric substances improve water retention in dryland biological soil crusts. Soil Biology and Biochemistry 116, 67–69. doi:10.1016/j.soilbio.2017.10.002

Ali, Sk.Z., Sandhya, V., Grover, M., Kishore, N., Rao, L.V., Venkateswarlu, B., 2009. Pseudomonas sp. strain AKM-P6 enhances tolerance of sorghum seedlings to elevated temperatures. Biology and Fertility of Soils 46, 45–55. doi:10.1007/s00374-009-0404-9

Amellal, N., Bartoli, F., Villemin, G., Talouizte, A., Heulin, T., 1999. Effects of inoculation of EPS-producing Pantoea agglomerans on wheat rhizosphere aggregation. Plant and Soil 211, 93–101. doi:10.1023/A:1004403009353

Amézketa, E., 1999. Soil Aggregate Stability: A Review. Journal of Sustainable Agriculture 14, 83–151. doi:10.1300/J064v14n02_08

Anderson-Teixeira, K.J., Davis, S.C., Masters, M.D., Delucia, E.H., 2009. Changes in soil organic carbon under biofuel crops. GCB Bioenergy 1, 75–96. doi:10.1111/j.1757-1707.2008.01001.x

Arnall, B., Jones, J., Pugh, B., Rocateli, A., Sanders, H., Warren, J., Zhang, H., 2018. Oklahoma Forage and Pasture Fertility Guide. Oklahoma State University, Stillwater, OK.

Bailey, V.L., Peacock, A.D., Smith, J.L., Bolton, H., 2002. Relationships between soil microbial biomass determined by chloroform fumigation–extraction, substrate-induced respiration, and phospholipid fatty acid analysis. Soil Biology and Biochemistry 34, 1385–1389. doi:10.1016/S0038-0717(02)00070-6

Benjamini, Y., Hochberg, Y., 1995. Controlling the False Discovery Rate: A Practical and Powerful Approach to Multiple Testing. Journal of the Royal Statistical Society: Series B (Methodological) 57, 289–300. doi:10.1111/j.2517-6161.1995.tb02031.x

Brejda, J.J., 2000. Fertilization of Native Warm-Season Grasses, in: Native Warm-Season Grasses: Research Trends and Issues, CSSA Special Publication. Crop Science Society of America and American Society of Agronomy, Madison, WI, pp. 177–200. doi:10.2135/cssaspecpub30.c12

Buyer, J.S., Sasser, M., 2012. High throughput phospholipid fatty acid analysis of soils. Applied Soil Ecology 61, 127–130. doi:10.1016/j.apsoil.2012.06.005

Caldwell, M.M., Dawson, T.E., Richards, J.H., 1998. Hydraulic lift: consequences of water efflux from the roots of plants. Oecologia 113, 151–161. doi:10.1007/s004420050363

Celik, G.Y., Aslim, B., Beyatli, Y., 2008. Characterization and production of the exopolysaccharide (EPS) from Pseudomonas aeruginosa G1 and Pseudomonas putida G12 strains. Carbohydrate Polymers 73, 178–182. doi:10.1016/j.carbpol.2007.11.021

Chang, W.-S., Mortel, M. van de, Nielsen, L., Guzman, G.N. de, Li, X., Halverson, L.J., 2007. Alginate Production by Pseudomonas putida Creates a Hydrated Microenvironment and Contributes to Biofilm Architecture and Stress Tolerance under Water-Limiting Conditions. Journal of Bacteriology 189, 8290–8299. doi:10.1128/JB.00727-07

Cheng, W., Gershenson, A., 2007. CHAPTER 2 - Carbon Fluxes in the Rhizosphere, in: Cardon, Z.G., Whitbeck, J.L. (Eds.), The Rhizosphere. Academic Press, Burlington, pp. 31–56. doi:10.1016/B978-012088775-0/50004-5

Chenu, C., 1995. Extracellular Polysaccharides: An Interface between Microorganisms and Soil Constituents, in: Huang, P.M., Berthelin, J., Bollag, J.-M., McGill, W.B. (Eds.), Environmental Impacts of Soil Component Interactions: Land Quality, Natural and Anthropogenic Organics, Volume I. Presented at the Impact of interactions of inorganic, organic, and microbiological soil components on environmental quality, Edmonton, Alberta, Canada, pp. 217–233.

Chenu, C., 1993. Clay-or sand-polysaccharide associations as models for the interface between micro-organisms and soil: water related properties and microstructure. Geoderma, International Workshop on Methods of Research on Soil Structure/Soil Biota Interrelationships 56, 143–156. doi:10.1016/0016-7061(93)90106-U

Chenu, C., Plante, A.T., 2006. Clay-sized organo-mineral complexes in a cultivation chronosequence: Revisiting the concept of the “primary organo-mineral complex.” European Journal of Soil Science 57, 596–607. doi:10.1111/j.1365-2389.2006.00834.x

Chenu, C., Roberson, E.B., 1996. Diffusion of glucose in microbial extracellular polysaccharide as affected by water potential. Soil Biology and Biochemistry 28, 877–884. doi:10.1016/0038-0717(96)00070-3

Cheshire, M.V., 1979. Nature and Origin of Carbohydrates in Soils. Academic Press, London, UK.

Cheshire, M.V., 1977. Origins and Stability of Soil Polysaccharide. Journal of Soil Science 28, 1–10. doi:10.1111/j.1365-2389.1977.tb02290.x

Chimento, C., Almagro, M., Amaducci, S., 2016. Carbon sequestration potential in perennial bioenergy crops: the importance of organic matter inputs and its physical protection. GCB Bioenergy 8, 111–121. doi:10.1111/gcbb.12232

Costa, O.Y.A., Raaijmakers, J.M., Kuramae, E.E., 2018. Microbial Extracellular Polymeric Substances: Ecological Function and Impact on Soil Aggregation. Frontiers in Microbiology 9. doi:10.3389/fmicb.2018.01636

D’Angelo, E., Crutchfield, J., Vandiviere, M., 2001. Rapid, Sensitive, Microscale Determination of Phosphate in Water and Soil. Journal of Environmental Quality 30, 2206–2209. doi:10.2134/jeq2001.2206

Dou, F.G., Hons, F.M., Ocumpaugh, W.R., Read, J.C., Hussey, M.A., Muir, J.P., 2013. Soil Organic Carbon Pools Under Switchgrass Grown as a Bioenergy Crop Compared to Other Conventional Crops. Pedosphere 23, 409–416. doi:10.1016/S1002-0160(13)60033-8

DuBois, Michel., Gilles, K.A., Hamilton, J.K., Rebers, P.A., Smith, Fred., 1956. Colorimetric Method for Determination of Sugars and Related Substances. Analytical Chemistry 28, 350–356. doi:10.1021/ac60111a017

Epskamp, S., 2019. semPlot: Path Diagrams and Visual Analysis of Various SEM Packages’ Output.

Fox, J., Weisberg, S., 2011. An R companion to applied regression, 2nd ed. ed. SAGE Publications, Thousand Oaks, Calif.

Frølund, B., Palmgren, R., Keiding, K., Nielsen, P.H., 1996. Extraction of extracellular polymers from activated sludge using a cation exchange resin. Water Research 30, 1749–1758. doi:10.1016/0043-1354(95)00323-1

Gelfand, I., Zenone, T., Jasrotia, P., Chen, J., Hamilton, S.K., Robertson, G.P., 2011. Carbon debt of Conservation Reserve Program (CRP) grasslands converted to bioenergy production. Proceedings of the National Academy of Sciences 108, 13864–13869. doi:10.1073/pnas.1017277108

Ghosh, S., Ghosh, P., Saha, P., Maiti, T.K., 2011. The extracellular polysaccharide produced by Rhizobium sp. isolated from the root nodules of Phaseolus mungo. Symbiosis 53, 75. doi:10.1007/s13199-011-0109-3

Gunina, A., Kuzyakov, Y., 2015. Sugars in soil and sweets for microorganisms: Review of origin, content, composition and fate. Soil Biology and Biochemistry 90, 87–100. doi:10.1016/j.soilbio.2015.07.021

Hall-Stoodley, L., Costerton, J.W., Stoodley, P., 2004. Bacterial biofilms: from the Natural environment to infectious diseases. Nature Reviews Microbiology 2, 95. doi:10.1038/nrmicro821

Idowu, O.J., 2003. Relationships Between Aggregate Stability and Selected Soil Properties in Humid Tropical Environment. Communications in Soil Science and Plant Analysis 34, 695–708. doi:10.1081/CSS-120018969

Jackson, R.B., Mooney, H.A., Schulze, E.-D., 1997. A global budget for fine root biomass, surface area, and nutrient contents. Proceedings of the National Academy of Sciences 94, 7362–7366. doi:10.1073/pnas.94.14.7362

Jastrow, J.D., Amonette, J.E., Bailey, V.L., 2007. Mechanisms controlling soil carbon turnover and their potential application for enhancing carbon sequestration. Climatic Change 80, 5–23. doi:10.1007/s10584-006-9178-3

Jastrow, J.D., Miller, R.M., Lussenhop, J., 1998. Contributions of interacting biological mechanisms to soil aggregate stabilization in restored prairie1The submitted manuscript has been created by the University of Chicago as operator of Argonne National Laboratory under Contract No. W-31-109-ENG-38 with the U.S. Department of Energy.1. Soil Biology and Biochemistry 30, 905–916. doi:10.1016/S0038-0717(97)00207-1

Jenkins, S., Swenson, T.L., Lau, R., Rocha, A.M., Aaring, A., Hazen, T.C., Chakraborty, R., Northen, T.R., 2017. Construction of Viable Soil Defined Media Using Quantitative Metabolomics Analysis of Soil Metabolites. Frontiers in Microbiology 8. doi:10.3389/fmicb.2017.02618

Jiao, Y., Cody, G.D., Harding, A.K., Wilmes, P., Schrenk, M., Wheeler, K.E., Banfield, J.F., Thelen, M.P., 2010. Characterization of extracellular polymeric substances from acidophilic microbial biofilms. Applied and Environmental Microbiology 76, 2916–2922. doi:10.1128/AEM.02289-09

Joergensen, R.G., Mueller, T., Wolters, V., 1996. Total carbohydrates of the soil microbial biomass in 0.5 M K2SO4 soil extracts. Soil Biology and Biochemistry 28, 1147–1153. doi:10.1016/0038-0717(96)00111-3

Kemper, W.D., Rosenau, R.C., 1986. Aggregate Stability and Size Distribution, in: Methods of Soil Analysis, Part 1. Physical and Mineralogical Methods (2nd Edition). pp. 425–442.

Kirkham, M.B., 2005. 19 - The Ascent of Water in Plants, in: Kirkham, M.B. (Ed.), Principles of Soil and Plant Water Relations. Academic Press, Burlington, pp. 315–340. doi:10.1016/B978-012409751-3/50019-0

Liang, C., Schimel, J.P., Jastrow, J.D., 2017. The importance of anabolism in microbial control over soil carbon storage. Nature Microbiology 2, 1–6. doi:10.1038/nmicrobiol.2017.105

Liao, J.D., Boutton, T.W., Jastrow, J.D., 2006. Organic matter turnover in soil physical fractions following woody plant invasion of grassland: Evidence from natural 13C and 15N. Soil Biology and Biochemistry, Ecosystems in Flux: Molecular and stable isotope Assessments of Soil Organic Matter Storage and Dynamics 38, 3197–3210. doi:10.1016/j.soilbio.2006.04.004

Ma, Z., Wood, C.W., Bransby, D.I., 2000. Soil management impacts on soil carbon sequestration by switchgrass. Biomass and Bioenergy 18, 469–477. doi:10.1016/S0961-9534(00)00013-1

Mao, Y., Li, X., Smyth, E.M., Yannarell, A.C., Mackie, R.I., 2014. Enrichment of specific bacterial and eukaryotic microbes in the rhizosphere of switchgrass (Panicum virgatum L.) through root exudates. Environmental Microbiology Reports 6, 293–306. doi:10.1111/1758-2229.12152

Marchus, K.A., Blankinship, J.C., Schimel, J.P., 2018. Environmental controls on extracellular polysaccharide accumulation in a California grassland soil. Soil Biology and Biochemistry 125, 86–92. doi:10.1016/j.soilbio.2018.07.009

Martin, J., 1946. Microorganisms and Soil Aggregation. Soil Science 61, 157–166.

McGowan, A.R., Min, D.-H., Williams, J.R., Rice, C.W., 2018. Impact of Nitrogen Application Rate on Switchgrass Yield, Production Costs, and Nitrous Oxide Emissions. Journal of Environmental Quality 47, 228–237. doi:10.2134/jeq2017.06.0226

McGowan, A.R., Nicoloso, R.S., Diop, H.E., Roozeboom, K.L., Rice, C.W., 2019. Soil Organic Carbon, Aggregation, and Microbial Community Structure in Annual and Perennial Biofuel Crops. Agronomy Journal 111, 128–142. doi:10.2134/agronj2018.04.0284

McLauchlan, K.K., Hobbie, S.E., Post, W.M., 2006. Conversion From Agriculture To Grassland Builds Soil Organic Matter On Decadal Timescales. Ecological Applications 16, 143–153. doi:10.1890/04-1650

Moffatt, H.H., 1973. Soil Survey of Caddo County, Oklahoma. United States Department of Agriculture Soil Conservation Service, Washington, D.C.

More, T.T., Yadav, J.S.S., Yan, S., Tyagi, R.D., Surampalli, R.Y., 2014. Extracellular polymeric substances of bacteria and their potential environmental applications. Journal of Environmental Management 144, 1–25. doi:10.1016/j.jenvman.2014.05.010

Nicolaus, B., Kambourova, M., Oner, E.T., 2010. Exopolysaccharides from extremophiles: From fundamentals to biotechnology. Environmental Technology 31, 1145–1158. doi:10.1080/09593330903552094

Oades, J.M., 1984. Soil organic matter and structural stability: mechanisms and implications for management. Plant and Soil 76, 319–337. doi:10.1007/BF02205590

O’Brien, S.L., Jastrow, J.D., 2013. Physical and chemical protection in hierarchical soil aggregates regulates soil carbon and nitrogen recovery in restored perennial grasslands. Soil Biology and Biochemistry 61, 1–13. doi:10.1016/j.soilbio.2013.01.031

Ontl, T.A., Cambardella, C.A., Schulte, L.A., Kolka, R.K., 2015. Factors influencing soil aggregation and particulate organic matter responses to bioenergy crops across a topographic gradient. Geoderma 255–256, 1–11. doi:10.1016/j.geoderma.2015.04.016

Ontl, T.A., Hofmockel, K.S., Cambardella, C.A., Schulte, L.A., Kolka, R.K., 2013. Topographic and soil influences on root productivity of three bioenergy cropping systems. New Phytologist 727–737. doi:10.1111/nph.12302@10.1002/(ISSN)1469-8137(CAT)VirtualIssues(VI)ScalingRootProcessesGlobalImpacts

Ophir, T., Gutnick, D.L., 1994. A Role for Exopolysaccharides in the Protection of Microorganisms from Desiccation. Appl. Environ. Microbiol. 60, 740–745.

Or, D., Smets, B.F., Wraith, J.M., Dechesne, A., Friedman, S.P., 2007. Physical constraints affecting bacterial habitats and activity in unsaturated porous media – a review. Advances in Water Resources, Biological processes in porous media: From the pore scale to the field 30, 1505–1527. doi:10.1016/j.advwatres.2006.05.025

Pal, A., Paul, A.K., 2013. Optimization of Cultural Conditions for Production of Extracellular Polymeric Substances (EPS) by Serpentine Rhizobacterium Cupriavidus pauculus KPS 201 [WWW Document]. Journal of Polymers. doi:10.1155/2013/692374

Parrish, D.J., Fike, J.H., 2005. The Biology and Agronomy of Switchgrass for Biofuels. Critical Reviews in Plant Sciences 24, 423–459. doi:10.1080/07352680500316433

Pedroso, G., De Ben, C., Hutmacher, R., Orloff, S., Putnam, D., Six, J., van Kessel, C., Wright, S., Linquist, B., 2011. Switchgrass is a promising, high-yielding crop for California biofuel. California Agriculture 65, 168–173.

Petersen, D.G., Blazewicz, S.J., Firestone, M., Herman, D.J., Turetsky, M., Waldrop, M., 2012. Abundance of microbial genes associated with nitrogen cycling as indices of biogeochemical process rates across a vegetation gradient in Alaska. Environmental Microbiology 14, 993–1008. doi:10.1111/j.1462-2920.2011.02679.x

Peterson, B.G., Carl, P., Boudt, K., Bennett, R., Ulrich, J., Zivot, E., Cornilly, D., Hung, E., Lestel, M., Balkissoon, K., Wuertz, D., 2018. PerformanceAnalytics: Econometric Tools for Performance and Risk Analysis.

Pett-Ridge, J., Firestone, M.K., 2017. Using stable isotopes to explore root-microbe-mineral interactions in soil. Rhizosphere, New Understanding of Rhizosphere Processes Enabled by Advances in Molecular and Spatially Resolved Techniques 3, 244–253. doi:10.1016/j.rhisph.2017.04.016

Quelas, J.I., López-García, S.L., Casabuono, A., Althabegoiti, M.J., Mongiardini, E.J., Pérez-Giménez, J., Couto, A., Lodeiro, A.R., 2006. Effects of N-starvation and C-source on Bradyrhizobium japonicum exopolysaccharide production and composition, and bacterial infectivity to soybean roots. Archives of Microbiology 186, 119–128. doi:10.1007/s00203-006-0127-3

R Core Team, 2018. R: A Language and Environment for Statistical Computing. R Foundation for Statistical Computing, Vienna, Austria.

Redmile-Gordon, M.A., Brookes, P.C., Evershed, R.P., Goulding, K.W.T., Hirsch, P.R., 2014. Measuring the soil-microbial interface: Extraction of extracellular polymeric substances (EPS) from soil biofilms. Soil Biology and Biochemistry 72, 163–171. doi:10.1016/j.soilbio.2014.01.025

Redmile-Gordon, M.A., Evershed, R.P., Hirsch, P.R., White, R.P., Goulding, K.W.T., 2015. Soil organic matter and the extracellular microbial matrix show contrasting responses to C and N availability. Soil Biology and Biochemistry 88, 257–267. doi:10.1016/j.soilbio.2015.05.025

Rinot, O., Levy, G.J., Steinberger, Y., Svoray, T., Eshel, G., 2019. Soil health assessment: A critical review of current methodologies and a proposed new approach. Science of The Total Environment 648, 1484–1491. doi:10.1016/j.scitotenv.2018.08.259

Roberson, E.B., 1991. Extracellular polysaccharide production by soil bacteria: environmental control and significance in agricultural soil. University of California, Berkeley.

Roberson, E.B., Firestone, M.K., 1992. Relationship between desiccation and exopolysaccharide production in a soil Pseudomonas sp. Applied and Environmental Microbiology 58, 1284–1291.

Roberson, E.B., Shennan, C., Firestone, M.K., Sarig, S., 1995. Nutritional Management of Microbial Polysaccharide Production and Aggregation in an Agricultural Soil. Soil Science Society of America Journal 59, 1587–1594. doi:10.2136/sssaj1995.03615995005900060012x

Robertson, G.P., Groffman, P.M., 2015. Chapter 14 - Nitrogen Transformations, in: Paul, E.A. (Ed.), Soil Microbiology, Ecology and Biochemistry (Fourth Edition). Academic Press, Boston, pp. 421–446. doi:10.1016/B978-0-12-415955-6.00014-1

Robertson, G.P., Hamilton, S.K., Barham, B.L., Dale, B.E., Izaurralde, R.C., Jackson, R.D., Landis, D.A., Swinton, S.M., Thelen, K.D., Tiedje, J.M., 2017. Cellulosic biofuel contributions to a sustainable energy future: Choices and outcomes. Science 356, eaal2324. doi:10.1126/science.aal2324

Rodrigues, R.R., Moon, J., Zhao, B., Williams, M.A., 2017. Microbial communities and diazotrophic activity differ in the root-zone of Alamo and Dacotah switchgrass feedstocks. GCB Bioenergy 9, 1057–1070. doi:10.1111/gcbb.12396

Rogers, S.L., Burns, R.G., 1994. Changes in aggregate stability, nutrient status, indigenous microbial populations, and seedling emergence, following inoculation of soil withNostoc muscorum. Biology and Fertility of Soils 18, 209–215. doi:10.1007/BF00647668

Rosseel, Y., 2012. lavaan: An R Package for Structural Equation Modeling. Journal of Statistical Software 48, 1–36.

Ruan, L., Bhardwaj, A.K., Hamilton, S.K., Robertson, G.P., 2016. Nitrogen fertilization challenges the climate benefit of cellulosic biofuels. Environmental Research Letters 11, 064007. doi:10.1088/1748-9326/11/6/064007

Russell, A.E., Kivlin, S.N., Hawkes, C.V., 2018. Tropical Tree Species Effects on Soil pH and Biotic Factors and the Consequences for Macroaggregate Dynamics. Forests 9, 184. doi:10.3390/f9040184

Sandhya, V., Ali, Sk.Z., 2015. The production of exopolysaccharide by Pseudomonas putida GAP-P45 under various abiotic stress conditions and its role in soil aggregation. Microbiology 84, 512–519. doi:10.1134/S0026261715040153

Schimel, J.P., 2018. Life in Dry Soils: Effects of Drought on Soil Microbial Communities and Processes. Annual Review of Ecology, Evolution, and Systematics 49, 409–432. doi:10.1146/annurev-ecolsys-110617-062614

Seki, K., 2007. SWRC fit – a nonlinear fitting program with a water retention curve for soils having unimodal and bimodal pore structure. Hydrology and Earth System Sciences Discussions 4, 407–437. doi:https://doi.org/10.5194/hessd-4-407-2007

Sheng, G.-P., Yu, H.-Q., Li, X.-Y., 2010. Extracellular polymeric substances (EPS) of microbial aggregates in biological wastewater treatment systems: A review. Biotechnology Advances 28, 882–894. doi:10.1016/j.biotechadv.2010.08.001

Sheng, G.P., Yu, H.Q., Yue, Z., 2006. Factors influencing the production of extracellular polymeric substances by Rhodopseudomonas acidophila. International Biodeterioration & Biodegradation, Surface Adhesion and Biotechnological Applications 58, 89–93. doi:10.1016/j.ibiod.2006.07.005

Shipley, B., 2016. Cause and correlation biology users guide path analysis structural equations and causal inference r 2nd edition | Ecology and co, 2nd ed. Cambridge university press.

Singer, M.J., Southard, R.J., Warrington, D.N., Janitzky, P., 1992. Stability of Synthetic Sand-Clay Aggregates after Wetting and Drying Cycles. Soil Science Society of America Journal 56, 1843–1848. doi:10.2136/sssaj1992.03615995005600060032x

Six, J., Conant, R.T., Paul, E.A., Paustian, K., 2002. Stabilization mechanisms of soil organic matter: Implications for C-saturation of soils. Plant and Soil 241, 155–176. doi:10.1023/A:1016125726789

Six, J., Elliott, E.T., Paustian, K., 2000. Soil macroaggregate turnover and microaggregate formation: a mechanism for C sequestration under no-tillage agriculture. Soil Biology and Biochemistry 32, 2099–2103. doi:10.1016/S0038-0717(00)00179-6

Six, J., Paustian, K., 2014. Aggregate-associated soil organic matter as an ecosystem property and a measurement tool. Soil Biology and Biochemistry 68, A4–A9. doi:10.1016/j.soilbio.2013.06.014

Sollins, P., Homann, P., Caldwell, B.A., 1996. Stabilization and destabilization of soil organic matter: mechanisms and controls. Geoderma 74, 65–105. doi:10.1016/S0016-7061(96)00036-5

Staudt, A.K., Wolfe, L.G., Shrout, J.D., 2012. Variations in exopolysaccharide production by Rhizobium tropici. Archives of Microbiology 194, 197–206. doi:10.1007/s00203-011-0742-5

Steinberger, R.E., Holden, P.A., 2004. Macromolecular composition of unsaturated *Pseudomonas aeruginosa* biofilms with time and carbon source. Biofilms 1, 37–47. doi:10.1017/S1479050503001066

Thomason, W.E., Raun, W.R., Johnson, G.V., Taliaferro, C.M., Freeman, K.W., Wynn, K.J., Mullen, R.W., 2005. Switchgrass Response to Harvest Frequency and Time and Rate of Applied Nitrogen. Journal of Plant Nutrition 27, 1199–1226. doi:10.1081/PLN-120038544

Tiemann, L.K., Grandy, A.S., 2015. Mechanisms of soil carbon accrual and storage in bioenergy cropping systems. GCB Bioenergy 7, 161–174. doi:10.1111/gcbb.12126

Tilman, D., Hill, J., Lehman, C., 2006. Carbon-Negative Biofuels from Low-Input High-Diversity Grassland Biomass. Science 314, 1598–1600. doi:10.1126/science.1133306

Tisdall, J.M., Oades, J.M., 1982. Organic matter and water-stable aggregates in soils. Journal of Soil Science 33, 141–163. doi:10.1111/j.1365-2389.1982.tb01755.x

Upadhyay, S., S Singh, J., Singh, D.P., 2011. Exopolysaccharide-Producing Plant Growth-Promoting Rhizobacteria Under Salinity Condition. Pedosphere 21, 214–222.

Venables, W.N., Ripley, B.D., 2002. Modern applied statistics with S, 4th ed. ed, Statistics and computing. Springer, New York.

Vogel, K.P., Brejda, J.J., Walters, D.T., Buxton, D.R., 2002. Switchgrass Biomass Production in the Midwest USA. Agronomy Journal 94, 413–420. doi:10.2134/agronj2002.0413

Wang, J., Yu, H.-Q., 2007. Biosynthesis of polyhydroxybutyrate (PHB) and extracellular polymeric substances (EPS) by Ralstonia eutropha ATCC 17699 in batch cultures. Applied Microbiology and Biotechnology 75, 871–878. doi:10.1007/s00253-007-0870-7

Wang, S., Redmile-Gordon, M., Mortimer, M., Cai, P., Wu, Y., Peacock, C.L., Gao, C., Huang, Q., 2019. Extraction of extracellular polymeric substances (EPS) from red soils (Ultisols). Soil Biology and Biochemistry 135, 283–285. doi:10.1016/j.soilbio.2019.05.014

Wickham, H., 2016. ggplot2: elegant graphics for data analysis, Second edition. ed, Use R! Springer, Cham.

Wolfaardt, G.M., Lawrence, J.R., Korber, D.R., 1999. Function of EPS, in: Wingender, J., Neu, T.R., Flemming, H.-C. (Eds.), Microbial Extracellular Polymeric Substances. Springer.

Yeats, T.H., Sorek, H., Wemmer, D.E., Somerville, C.R., 2016. Cellulose Deficiency Is Enhanced on Hyper Accumulation of Sucrose by a H+-Coupled Sucrose Symporter. Plant Physiology 171, 110–124. doi:10.1104/pp.16.00302

Zhalnina, K., Louie, K.B., Hao, Z., Mansoori, N., Rocha, U.N. da, Shi, S., Cho, H., Karaoz, U., Loqué, D., Bowen, B.P., Firestone, M.K., Northen, T.R., Brodie, E.L., 2018. Dynamic root exudate chemistry and microbial substrate preferences drive patterns in rhizosphere microbial community assembly. Nature Microbiology 3, 470. doi:10.1038/s41564-018-0129-3

Zhang, Q., Yan, T., 2012. Correlation of Intracellular Trehalose Concentration with Desiccation Resistance of Soil Escherichia coli Populations. Appl. Environ. Microbiol. 78, 7407–7413. doi:10.1128/AEM.01904-12

Zhu, S., Vivanco, J.M., Manter, D.K., 2016. Nitrogen fertilizer rate affects root exudation, the rhizosphere microbiome and nitrogen-use-efficiency of maize. Applied Soil Ecology 107, 324–333. doi:10.1016/j.apsoil.2016.07.009

Zuur, A.F., Ieno, E.N., Elphick, C.S., 2010. A protocol for data exploration to avoid common statistical problems. Methods in Ecology and Evolution 1, 3–14. doi:10.1111/j.2041-210X.2009.00001.x

