## SUPPLEMENTAL METHODS; SUPPLEMENTAL TABLES S1-S3; SUPPLEMENTAL FIGURES S1-S5 for "Microbial extracellular polysaccharide production and aggregate stability controlled by Switchgrass (*Panicum virgatum*) root biomass and soil water potential"

### SUPPLEMENTAL MATERIALS

### SUPPLEMENTAL METHODS

#### Labeling setup

All chambers under a labeling regime were connected to a single continuous gas line by solenoids, with gas flow into each individual chamber controlled via the CR1000 data logger (**Fig. 1B**). Labeling chambers were created by affixing another clear, impact-resistant polycarbonate tube (122 cm x 19.7 cm) to the top of each mesocosm and around the growing SG using PAR-transmissive gas-tight tape (LI-COR, Lincoln, NE). These labeling chambers were themselves capped with PAR-transmissive film (McMaster-Carr, Elmhurst, IL) affixed with the same tape. $CO_2$ was delivered to each chamber under the two labeling regimes dynamically throughout the day to account for increased photosynthetic activity throughout the morning and early afternoon, before tapering off in the late afternoon and evening. Overnight, $CO_2$ delivery was halted and [$CO_2$] was allowed to build up in the chamber headspace. In the morning, $CO_2$ delivery would only begin once the chamber headspace [$CO_2$] was reduced below 400 ppm. Thus, both labeling regimes can be considered to have been exposed to elevated $CO_2$ during and immediately after nighttime accumulation of respired soil $CO_2$.

### SUPPLEMENTAL TABLES

**Table S1:** Multiple linear regression models describing relations between most highly explanatory soil factors and water stable aggregates, after removing collinear explanatory soil factors with VIF above 3 (Zuur et al., 2010).

| Factor | | | Soil water potential* | | pH | |
|---|---|---|---|---|---|---|
| | Model $R^2$ | Model P | β[§] | P | β | P |
| | **0.393** | **0.0005798** | **0.03749** | **0.00936** | **-0.12785** | **0.00353** |

* Absolute value soil water potential units

[§] Factor specific slope when other factors are constant

**Table S2:** Soil chemistry and SG root biomass of bulk soil recovered from each treatment and horizon at the end of the study, $^{13}C$ enriched EPS and total soil C measured in the surface horizon at the end of the study, soil chemistry and texture of soil from initial soil horizons, and initial field bulk density of each horizon. There were no significant differences in total carbon, nitrogen, or phosphorus between treatments. There were also no significant differences in total phosphorus between horizons, but there were significant differences observed in total carbon and nitrogen.

| | | | Treatment | | | | |
|---|---|---|---|---|---|---|---|
| Element | Horizon | Initial | Control | N | NP | P | W |
| EPS | A | | 10.3±1.5 | 12.0±0.5 | 14.6±1.6 | 10.9±2.3 | 11.0±0.5 |
| (µg glucose equivalent | B | | 7.05±1.08 | 7.29±1.50 | 7.66±1.75 | 6.48±1.91 | 6.50±1.85 |
| $g^{-1}$ dry soil) | C | | 2.91±1.43 | 4.25±1.77 | 3.32±0.85 | 2.67±1.11 | 3.24±1.71 |
| Root biomass | A | | 4.15±0.50 | 4.13±0.99 | 6.07±1.02 | 4.83±0.90 | 3.23±0.62 |
| (g dry root $g^{-1}$ dry soil) | B | | 1.92±0.43 | 2.94±0.32 | 2.79±0.60 | 2.62±0.14 | 1.67±0.29 |
| | C | | 1.64±0.39 | 1.90±0.36 | 1.94±0.51 | 1.96±0.24 | 1.57±0.25 |
| pH | A | | 5.27±0.12 | 4.94±0.13 | 4.96±0.14 | 5.31±0.15 | 5.33±0.07 |
| | B | | 6.20±0.08 | 6.25±0.09 | 6.25±0.04 | 6.11±0.07 | 6.09±0.07 |
| | C | | 6.47±0.16 | 6.46±0.08 | 6.43±0.07 | 6.40±0.12 | 6.40±0.13 |
| Soil moisture | A | | 527±352 | 2360±1380 | 11100±15700 | 935±680 | 12100±5700 |
| (-kPa) | B | | 1040±910 | 4690±1180 | 6040±2250 | 1840±1400 | 7570±2940 |
| | C | | 571±388 | 2040±690 | 2330±270 | 810±322 | 2290±690 |
| DOC | A | | 26.5±1.6 | 30.3±1.8 | 32.4±1.8 | 25.7±2.1 | 24.9±3.5 |
| (µg $g^{-1}$) | B | | 17.9±2.6 | 17.9±1.6 | 22.0±4.6 | 18.8±2.6 | 21.7±0.9 |

| | | | | | | | |
|---|---|---|---|---|---|---|---|
| | C | | 17.2±2.2 | 16.7±1.5 | 17.0±2.7 | 17.7±5.0 | 18.6±2.8 |
| DN | A | | 3.20±0.18 | 21.4±3.3 | 12.3±3.0 | 3.43±0.21 | 3.29±0.38 |
| (μg $g^{-1}$) | B | | 2.46±1.96 | 9.48±5.87 | 7.02±1.15 | 1.96±0.43 | 2.41±0.44 |
| | C | | 1.35±0.64 | 1.40±0.32 | 2.56±0.54 | 1.46±0.84 | 1.49±0.45 |
| EPS $^{13}C$ (ug glucose equivalent $g^{-1}$ dry soil) | A | | 0.00721±0.00047 | 0.00891±0.00106 | 0.00503±0.00123 | 0.00588±0.0035 | 0.00294±0.00100 |
| Total carbon $^{13}C$ | A | | 7.24±4.16 | 7.25±6.03 | 2.18±0.99 | 2.92±0.87 | 2.35±0.47 |
| Total carbon | A | 0.390±0.003 | 0.368±0.023 | 0.377±0.025 | 0.414±0.026 | 0.372±0.038 | 0.378±0.045 |
| (%) | B | 0.203±0.018 | 0.214±0.080 | 0.174±0.014 | 0.174±0.019 | 0.176±0.016 | 0.176±0.013 |
| | C | 0.106±0.005 | 0.104±0.012 | 0.107±0.005 | 0.112±0.010 | 0.104±0.009 | 0.108±0.008 |
| Total nitrogen | A | 0.036±0. | 0.0320±0.0130 | 0.0313±0.0097 | 0.0320±0.0062 | 0.0290±0.0077 | 0.0318±0.0091 |
| (%) | B | 0.014±0. | 0.0143±0.0106 | 0.00900±0.00510 | 0.0103±0.0051 | 0.00950±0.00509 | 0.00983±0.00417 |
| | C | 0.006±0. | 0.00283±0.00371 | 0.00233±0.00301 | 0.00317±0.00319 | 0.00200±0.00276 | 0.00250±0.00351 |
| Total phosphorus | A | 5.66±0.44 | 4.69±0.67 | 4.64±0.57 | 5.69±0.91 | 4.89±0.97 | 4.62±0.54 |
| (ppm) | B | 4.19±0.27 | 3.52±0.33 | 3.42±0.31 | 3.81±0.81 | 3.62±0.27 | 3.49±0.21 |
| | C | 5.63±0.95 | 4.44±0.24 | 5.00±0.66 | 4.99±0.48 | 4.78±0.45 | 4.58±0.39 |
| Clay (%) | A | 14.3±2.9 | | | | | |
| | B | 11.4±0.0 | | | | | |
| | C | 13.4±1.7 | | | | | |
| Silt (%) | A | 17.1±5.7 | | | | | |

| | | |
|---|---|---|
| | B | 11.4±2.9 |
| | C | 9.51±1.65 |
| Sand (%) | A | 68.6±4.9 |
| | B | 77.2±2.8 |
| | C | 78.1±1.7 |
| Bulk density | A | 1.41±0.04 |
| (g dry soil $cm^{-3}$) | B | 1.53±0.18 |
| | C | 1.64±0.07 |

**Table S3:** Hexose to pentose ratios of polysaccharides extracted from soil, in the greenhouse experiment and from Red River site, and from Switchgrass roots, from Red River site.

| | (G+M)/(A+X) | M/(A+X) |
|---|---|---|
| Control | 4.19±0.18 | 2.27±0.10 |
| N | 3.86±0.13 | 2.23±0.10 |
| NP | 3.77±0.20 | 2.19±0.14 |
| P | 4.08±0.06 | 2.36±0.07 |
| Low water | 3.72±0.22 | 2.18±0.15 |
| RR- root | 1.45±0.17 | 0.55±0.11 |
| RR- soil | 5.35±1.08 | 2.77±0.67 |

### SUPPLEMENTAL FIGURES

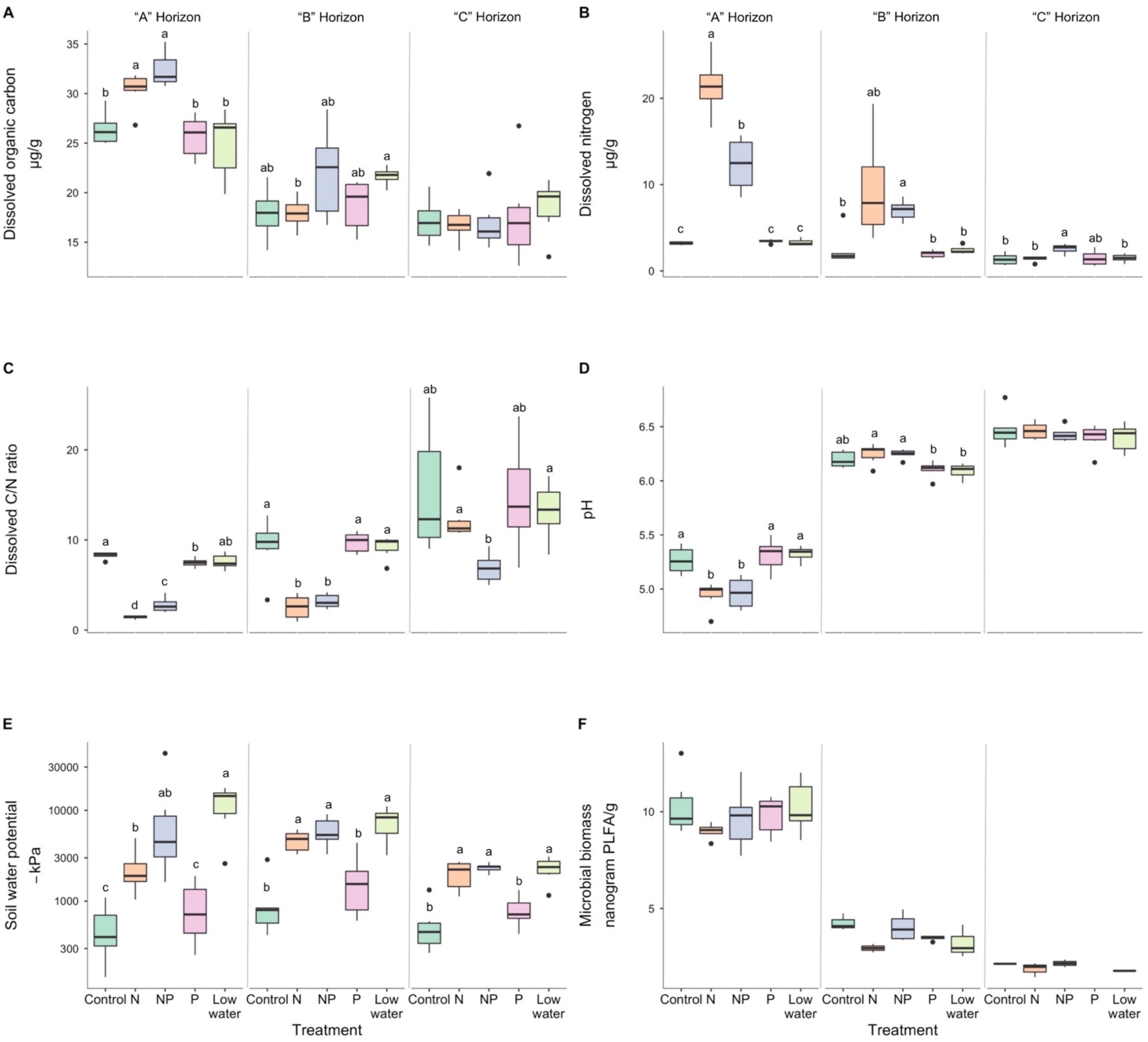


**Figure S1.** Differences between soil factors across treatments and soil horizons. Box-whisker plots of A) dissolved organic carbon (micrograms C per gram dry soil), B) dissolved nitrogen (micrograms N per gram dry soil), C) the ratio of dissolved organic C to dissolved N, D) soil pH, E) soil water potential (negative kilopascals) and F) microbial biomass (nanograms of phospholipid fatty acids per gram dry soil) recovered from bulk soil by treatment and horizon. Lowercase letters indicate significant differences between treatments (Welch's t-test $P < 0.05$, after Benjamini-Hochberg correction for multiple testing).

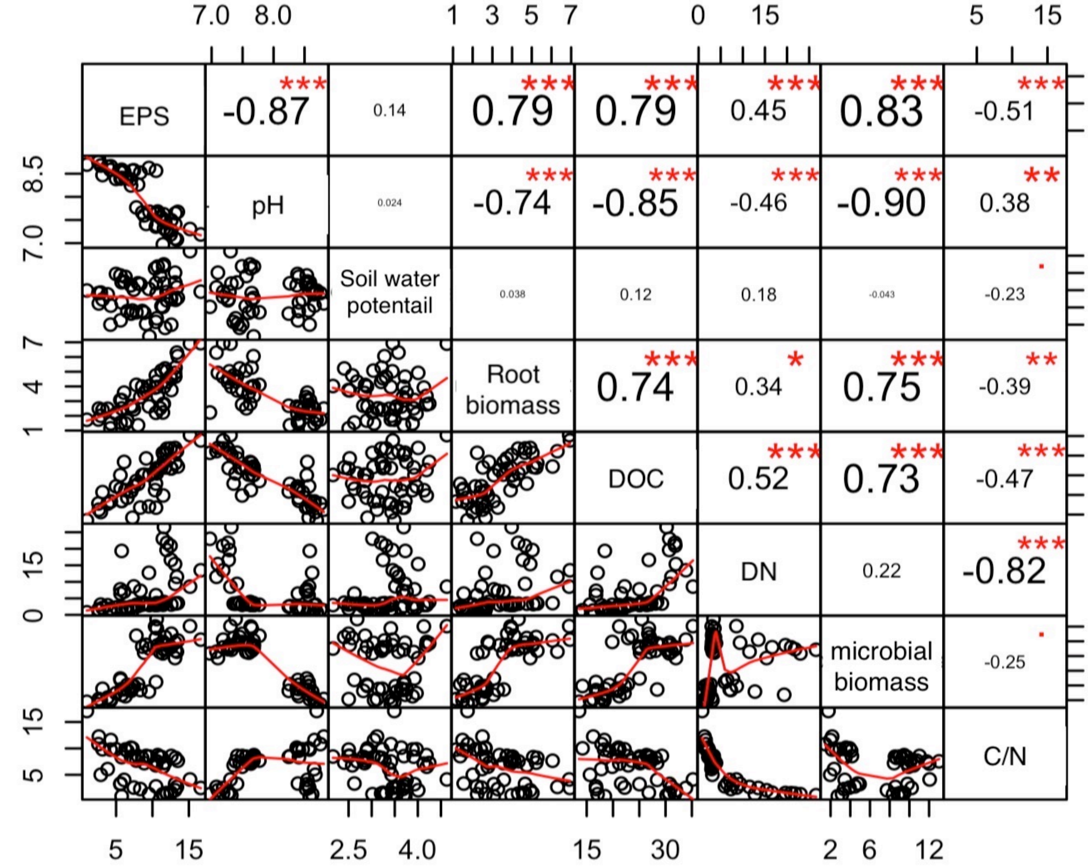


**Figure S2.** Matrix of correlations between measured soil characteristics, across all horizons and treatments. An intersection of two variables on the bottom left contains a scatterplot of their relationship; intersections on the top right contain corresponding Spearman correlation coefficients for these relationships. Red asterisks denote significant correlations ($P < 0.05$ for *, $< 0.01$ for **, and $< 0.001$ for ***).

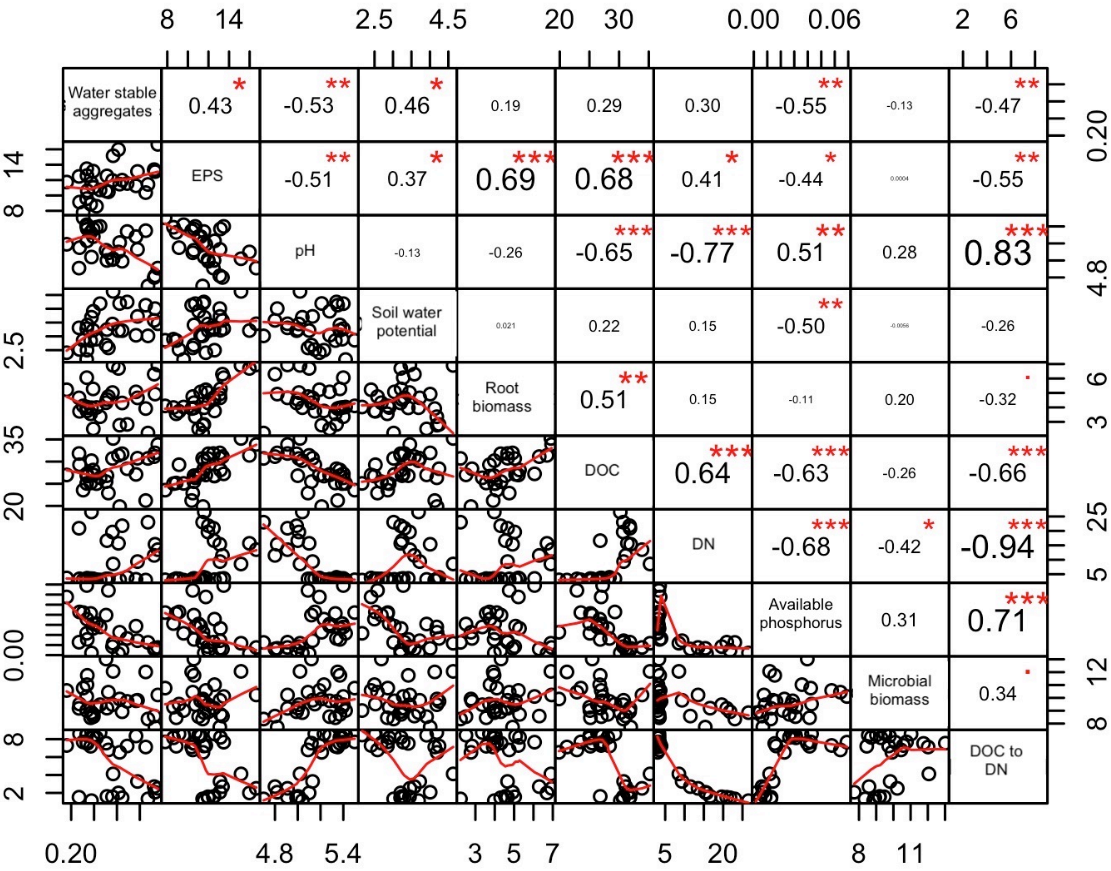


**Figure S3.** Matrix of correlations between measured soil characteristics, across all treatments within the A horizon only. An intersection of two variables on the bottom left contains a scatterplot of their relationship; intersections on the top right contain corresponding Spearman correlation coefficients for these relationships. Red asterisks denote significant correlations (P < 0.05 for *, < 0.01 for **, and < 0.001 for ***).

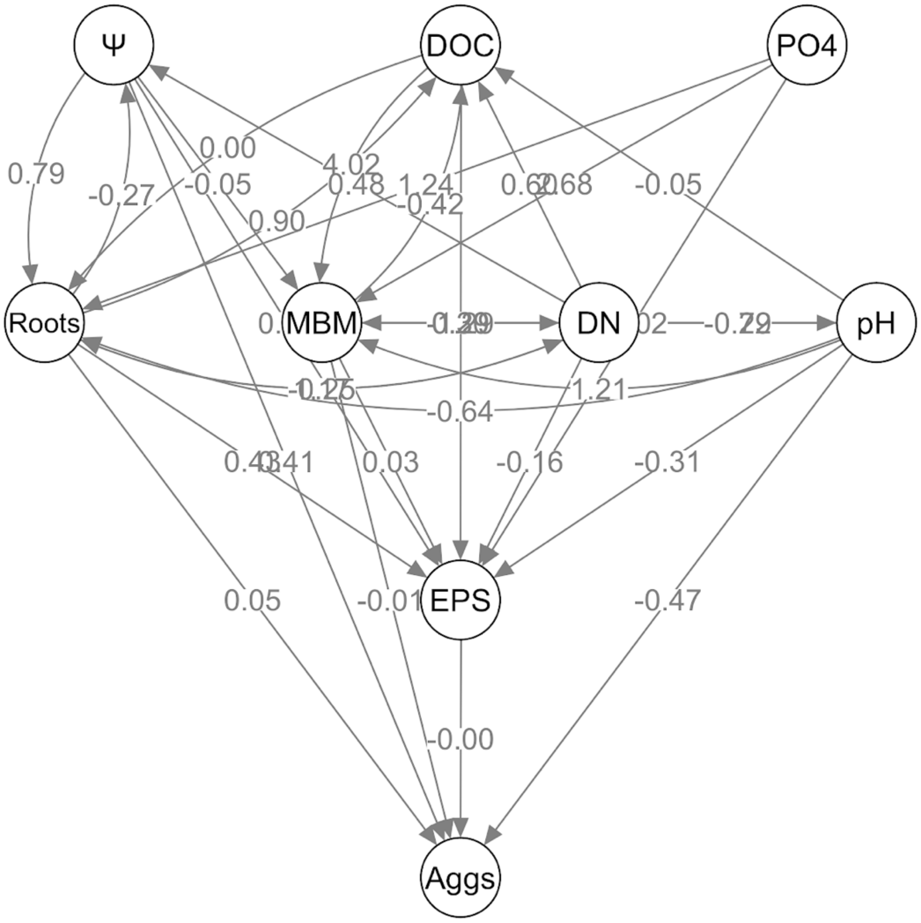


**Figure S4.** Path analysis of soil factors eventually affecting EPS and soil aggregate stability. Full, un-constrained model for initial framing of path analysis based on theoretical framework of interactions between the relevant measured variables, before non-significant edges were iteratively removed. Node labels correspond to the following measured variables: EPS content (EPS), frequency of water-stable aggregates (Aggs), soil water potential (Ψ), pH, SG root biomass (Roots), dissolved organic carbon (DOC), total dissolved nitrogen (DN), microbial biomass measured by PLFA (MBM), and phosphate accumulation on anion exchange membranes over the course of the study (PO4).

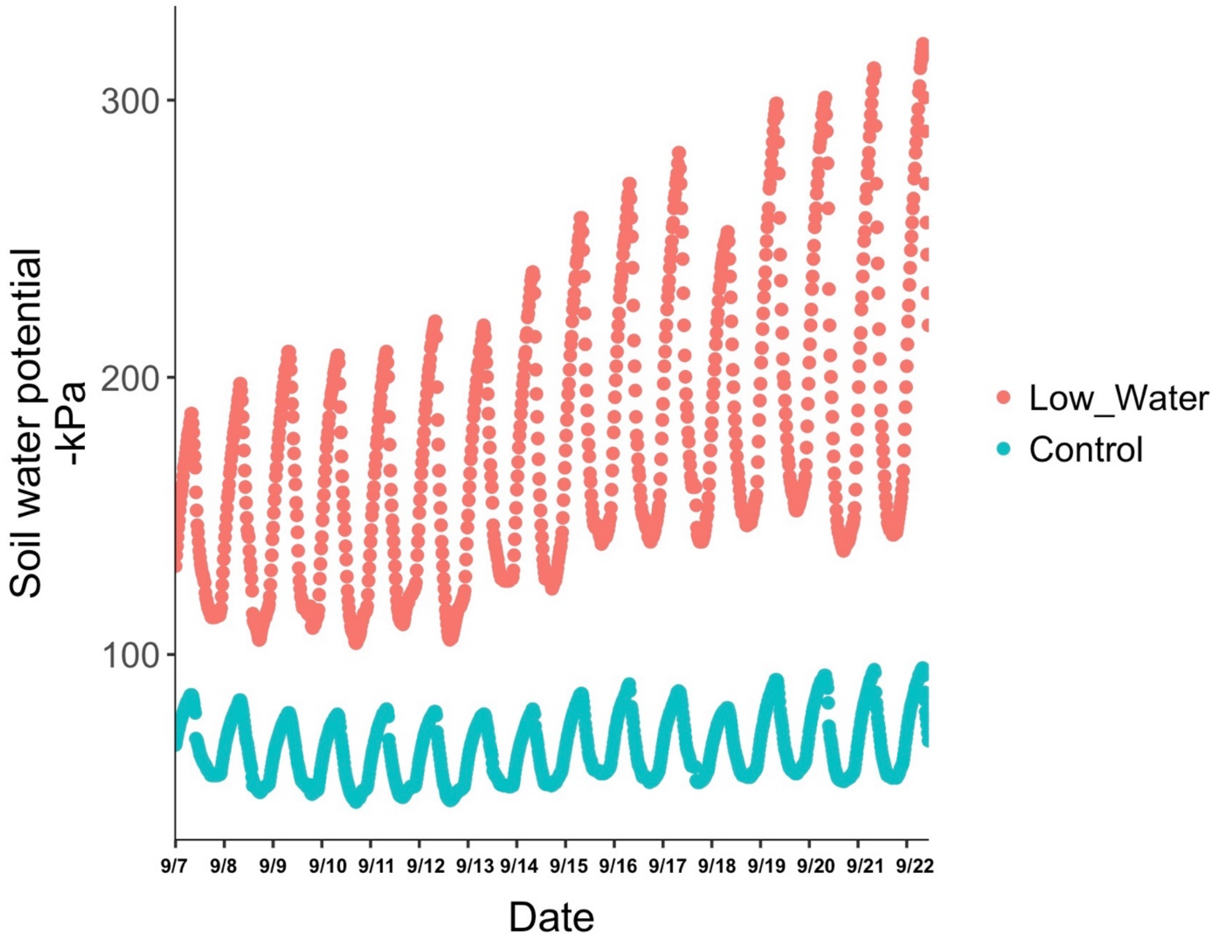


**Figure S5.** Daily changes in soil water potential during two weeks (including 12 days of labeling) before the destructive harvest from the A horizon of four mesocosms, two from the low water treatment and two from the control treatment. Each dot represents a measurement taken every 20 minutes.
